# Identifying Plasmodium falciparum transmission patterns through parasite prevalence and entomological inoculation rate

**DOI:** 10.1101/2021.01.14.426709

**Authors:** Benjamin Amoah, Robert S. McCann, Alinune N. Kabaghe, Monicah Mburu, Michael G. Chipeta, Paula Moraga, Steven Gowelo, Tinashe Tizifa, Henk van den Berg, Themba Mzilahowa, Willem Takken, Michéle van Vugt, Kamija S. Phiri, Peter J. Diggle, Dianne J. Terlouw, Emanuele Giorgi

## Abstract

Monitoring malaria transmission is a critical component of efforts to achieve targets for elimination and eradication. Two commonly monitored metrics of transmission intensity are parasite prevalence (PR) and the entomological inoculation rate (EIR). Using geostatistical methods, we investigate the relationship between *Plasmodium falciparum* PR and EIR using data collected over 38 months in a rural area of Malawi. Our results indicate that hotspots identified through the EIR and PR partly overlapped during high transmission seasons but not during low transmission seasons. The estimated relationship showed a one-month delayed effect of EIR on PR such that at low transmission levels increases in EIR are associated with rapid rise in PR, but at high transmission levels, decreases in EIR do not translate into notable reductions in PR. Our study emphasises the need for integrated malaria control strategies that combines vector and human host managements monitored by both entomological and parasitaemia indices.

## Introduction

National malaria control programmes, working in collaboration with global stakeholders, have achieved extensive intervention coverage over the last two decades, leading to significant reductions in morbidity and mortality due to malaria (***Bhatt et al., 2015b***). However, malaria is still a leading global health problem. The previous successes and current challenges have motivated ambitious, yet feasible, global and national targets towards malaria elimination. A key component of efforts to achieve these targets is surveillance and monitoring, which is critical for continued assessment of intervention effectiveness, identification of areas or groups at the highest risk, and guiding the development and implementation of new intervention strategies (***World Health Organization, 2015***).

A wide range of metrics exists for monitoring malaria parasite transmission. The strengths and limitations of each metric are related, in part, to the step of the parasite transmission cycle it measures (***Tusting et al., 2014***). These strengths and weaknesses, including the sensitivity of each metric, which vary across epidemiological settings and as parasite transmission declines within a given setting (***The malERA Refresh Consultative Panel on Characterising the Reservoir and Measuring Transmission, 2017***). Two of the most commonly monitored metrics are the prevalence of *Plasmodium* parasites and the entomological inoculation rate (EIR), especially in moderate to high transmission settings.

The prevalence of *Plasmodium* parasites in the human population at a given time point (i.e. the parasite rate; PR) approximates the reservoir of hosts potentially available to transmit the parasite from humans to mosquitoes. Whereas only the gametocyte stage of the parasite contributes to transmission, it remains relatively expensive to detect this stage of the parasite. Whereas rapid diagnostic tests (RDTs) that primarily detect asexual-stage antigens are inexpensive and easily deployed in large-scale community-based surveys (***Poti et al., 2020***), their limit of detection (50-200 parasites/µl) is higher than that of expert microscopy or PCR (***Chiodini, 2014***), so that RDT-based estimates of PR are biased by excluding low-density infections. Despite these limitations, RDT-based cross-sectional surveys to measure PR capture both symptomatic and asymptomatic infections, which is important because both are likely to contribute to transmission (***Bousema et al., 2014***; ***Slater et al., 2019***), and changes in PR over time can indicate changes in transmission.

EIR provides an estimate of the intensity of parasite transmission from mosquitoes to humans, expressed as the number of infectious bites received per person per unit time. EIR is calculated by multiplying the number of malaria vector bites per person per unit time, also known as the human biting rate (HBR), by the proportion of vectors carrying the infectious sporozoite stage of malaria parasites, referred to as the sporozoite rate (SR) (***Onori and Grab, 1980***). The accuracy and precision of EIR estimates, therefore, depends on the accuracy and precision with which HBR and SR can be measured (***Tusting et al., 2014***). Two common methods for measuring HBR are the human landing catch and the Centers for Disease Control and Prevention Light Trap, but inter-individual variation in attractiveness to mosquitoes restricts standardisation across sampling points for both of these methods (***Knols et al., 1995***; ***Qiu et al., 2006***). Alternative methods for estimating HBR include the Suna trap, which uses a synthetic blend of volatiles found on human skin and carbon dioxide to attract host-seeking *Anopheles* mosquitoes (***Mukabana et al., 2012***; ***Menger et al., 2014***; ***Hiscox et al., 2014***). The standardised odour blend allows for reliable comparisons among trapping locations (***Mburu et al., 2019***). Regardless of the method used to estimate HBR, the precision of SR decreases as the number of mosquitoes collected decreases. Despite these limitations, EIR is a vital metric of malaria parasite transmission because it directly describes human exposure to malaria parasites before post-inoculation factors such as immunity, nutrition, and access to health care (***Killeen et al., 2000***). Moreover, EIR provides information about the relative contributions of different vector species to transmission, which can impact malaria intervention effectiveness based on interspecies differences in biting behaviours related to time and location, non-human blood-meal hosts, larval ecology, and insecticide resistance profiles (***Ferguson et al., 2010***).

Malaria parasite transmission is heterogeneous in space and time at fine resolution due to several factors, including the availability of larval mosquito habitat, socioeconomics, human behaviour and genetics, and malaria intervention coverage (***Carter et al., 2000***; ***Bousema et al., 2012***; ***McCann et al., 2017b***). Repeated cross-sectional surveys continousely carried out in communities can reveal this fine-resolution heterogeneity (***Roca-Feltrer et al., 2012***), providing timely estimates of malaria control progress at the sub-district level and potentially identifying hotspots of malaria parasite transmission for targeted intervention (***Kabaghe et al., 2017***; ***Bousema et al., 2016***). However, understanding this heterogeneity and identifying hotspots in a way that is meaningful for control programmes remains challenging (***Stresman et al., 2019***), in part because hotspot location and size can depend on which metric is used (***Stresman et al., 2017***). Given that PR and EIR are indicative of components of the parasite transmission cycle that are separated by multiple complex steps, each metric provides partial but useful information about the underlying risk of transmission. Therefore, measuring and mapping both metrics can provide a fuller picture of parasite transmission (***Cohen et al., 2017***).

Additionally, modelling the functional relationship between EIR and PR can provide further insights into the underlying malaria epidemiology. For example, the functional relationship between EIR and PR can then, for monitoring and evaluation purposes, be used to quantify the average reductions in prevalence that may be gained as a result of reductions in EIR, and conversely, the expected increase in prevalence when the number of infectious mosquito bites increases. Previous studies have modelled the functional relationship between EIR and PR using paired estimates of EIR and PR from sites representing a wide range of EIR and PR in Africa (***Beier et al., 1999***; ***Smith et al., 2005***). These meta-analyses used one estimate each of EIR and PR per site from studies conducted before 2004 and excluded sites with reported malaria control activities, revealing consistently high PR (above 50%) for sites with EIR greater than 15 infectious bites (ib)/person/year and a steep decrease in PR with decreasing EIR when EIR is below 15 ib/person/year. In the current study, we investigate the EIR-PR relationship over a finer time resolution of one month for 38 months, within a single geographical region. The EIR-PR relationship in this context, therefore, takes into account subannual changes in transmission, likely driven by seasonal weather patterns, and other (year-to-year) spatiotemporal variabilities, likely driven by a combination of climatic variation and changes in malaria control activities.

The joint monitoring of EIR and PR in space and time allows us, in this paper, to investigate and find answers to the following questions. (1) How do spatiotemporal patterns of EIR and PR compare? (2) Do EIR and PR lead to the identification of the same malaria hotspots? (3) How do changes in EIR affect PR at different transmission levels? (4) Does EIR have a lagged effect on PR? (5) Does the EIR-PR relationship vary between women of reproductive age and children between 6 and 60 months of age? To answer these questions, we first map *P. falciparum* entomological inoculation rate (PfEIR) and *P. falciparum* parasite prevalence (PfPR) at a high spatiotemporal resolution to identify and compare their spatial heterogeneities and temporal patterns. We then consider several statistical models for the relationship between PfEIR and PfPR, which can be distinguished as follows: mechanistic models that are based on different epidemiological assumptions and empirical models where the data inform the PfEIR-PfPR relationship. Finally, we discuss the possible implications of the estimated relationships for monitoring malaria control strategies.

## Methods

### Study site

This study was part of the Majete Malaria Project (MMP), an integrated malaria control project in Chikwawa District, Malawi. The catchment area of MMP consisted of three distinct geographical regions, referred to as Focal Areas A, B and C (***Figure 1***), with a total population of about 25,000 people living in 6,600 households in 65 villages.

**Figure 1.**
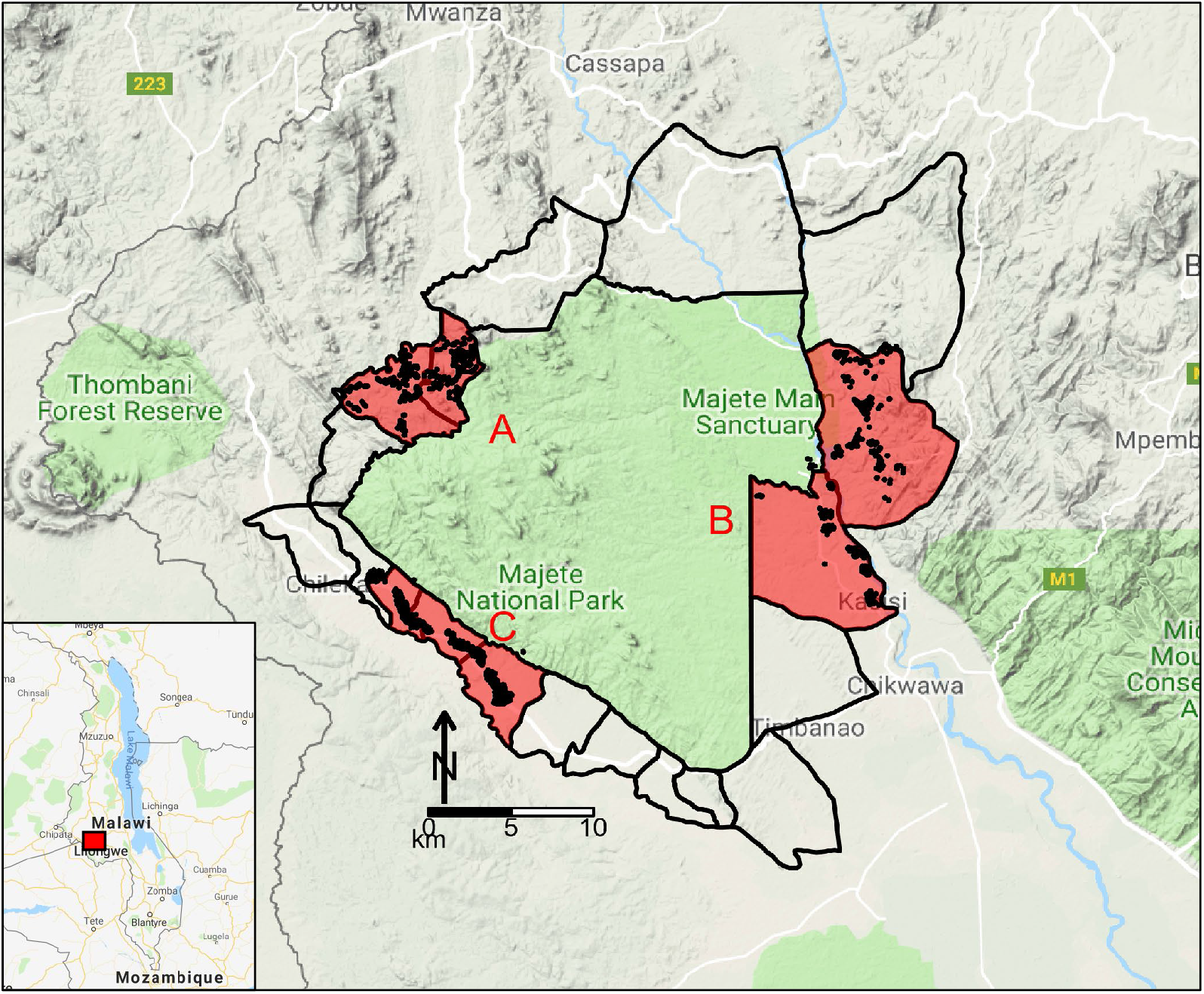
Map of study site. Map of Malawi (insert) highlighting the Majete Wildlife Reserve and the borders of 19 community-based organisations (CBOs) surrounding the Majete perimeter. Three focal areas (red patches), labelled as A, B, and C, show the households (black points) selected for the parasitaemia and entomological surveys by the Majete Malaria Project (MMP).

Chikwawa experiences highly variable rainfall during its single rainy season, which spans November/December to April/May. Temperatures are generally high, with daily maximum temperatures in December averaging 37.6 °C, and in July averaging 27.6 °C (***Joshua et al., 2016***). During the rainy season, the Shire and Mwanza rivers, which run near the study area, create marshy habitats, paddies, occasional depressions and watering holes, suitable as larval habitats for *Anopheles funestus s*.*s*., *Anopheles arabiensis* and *Anopheles gambiae s*.*s*. (***Spiers et al., 2002***). Dry season larval habitats consist primarily of burrow pits and pools of standing water along seasonal stream beds.

Malaria control in the district is implemented through the Chikwawa District Health Office. During the study period, interventions applied throughout the study area included the continuous provision of insecticide-treated nets (ITNs) to pregnant women and children under five years old, mass distribution campaigns of ITNs, intermittent preventative therapy for pregnant women, and malaria case diagnosis and treatment with artemisinin-based combination therapy. The only mass distribution of ITNs in the district during the study period occurred in April 2016. As part of the MMP, a randomised trial was conducted to assess the effectiveness of additional, community-implemented malaria interventions between May 2016 and May 2018 (***McCann et al., 2017a***). The trial interventions were implemented at the village level, with villages assigned to one of four groups: a) no additional interventions; b) larval source management; c) house improvement; and d) both larval source management and house improvement (***McCann et al., 2017a***; ***van den Berg et al., 2018***).

### Data

To quantify PfPR and PfEIR over the course of the study, a rolling malaria indicator survey (rMIS) was conducted in conjunction with mosquito sampling (***Roca-Feltrer et al., 2012***). In the first two rounds of baseline data collection (April through August 2015), an inhibitory geostatistical sampling design (IGSD) was used to select 300 and 270 households, respectively, for the rMIS from an enumeration database of all households in the catchment area (***Chipeta et al., 2017***). The IGSD helped to ensure that randomly sampled households are relatively uniformly spaced over the study region by requiring each pair of sampled households to be separated by a distance of at least 0.1km, which increases the efficiency of hotspot detection (***Kabaghe et al., 2017***). In the three subsequent rounds of data collection during the baseline, an adaptive geostatistical sampling design (AGSD) was used to select 270 households per round (***Chipeta et al., 2016***). With AGSD, new households for the current round of rMIS were chosen from regions with high standard errors of estimated prevalence, based on data from all previous rounds. In the baseline period, previously sampled households were not eligible for sampling in subsequent rounds. For the trial period (starting May 2016), IGSD was again used to select households from the enumeration database of all households. All households were eligible for selection in each round of the trial period regardless of whether they were selected in a previous round. At each round of rMIS data collection in the baseline and trial phases, respectively, 75% and 72% of the households chosen at each round of the rMIS were then randomly selected for mosquito sampling.

In each sampled household, children under five (0.5-5 y/o) and women of reproductive age (15-49 y/o) were tested for *P. falciparum* using an RDT (SD BIOLINE Malaria Ag P.f. HRP-II, Standard Diagnostics, Yongin-si, Republic of Korea).

Mosquitoes were sampled from 5pm to 7am using Suna traps (Biogents AG, Regensburg, Germany) (***Hiscox et al., 2014***) with MB5 blend plus *co*_2_ to mimic human odour (***Mburu et al., 2019***). For a selected household in a surveillance round, the trap was set for one night indoors and one night outdoors, with the order of indoor/outdoor determined randomly. For households where the residents were sleeping in more than one building, a trap was set at each building. Trapped female anophelines were preserved using a desiccant and identified using standard morphological and molecular techniques (***Gillies and Coetzee, 1987***; ***Koekemoer et al., 2002***; ***Scott et al., 1993***). Female anophelines were further tested for the presence of *P. falciparum* in their head and thorax, after removing the abdomen, using quantitative polymerase chain reaction (qPCR) (***Bass et al., 2008***; ***Perandin et al., 2004***). Specimens with a Ct value below 37.0 were considered positive for *P. falciparum*.

### Environmental and climatic factors

Environmental and climatic factors affect the abundance and suitability of water bodies that support the development of immature mosquitoes (***Madder et al., 1983***; ***Loetti et al., 2011***), the duration of mosquito development (***Ciota et al., 2014***; ***Loetti et al., 2011***; ***Craig et al., 1999***), mosquito host-seeking and biting behaviour, and the development rate of malaria parasites in mosquitoes (***Rumisha et al., 2014***; ***Amek et al., 2011***).

Using hourly measurements of temperature and relative humidity (RH) from a weather station in each focal area, we computed the average temperature and RH for different ranges of days before the day of data collection (***Appendix 1***).

Spectral indices, namely normalised difference vegetation index (NDVI) and enhanced vegetation index (EVI), were computed using remotely sensed multi-spectral imagery from the Landsat 8 satellite. These data are freely available from the United States Geological Survey (USGS) Earth Explorer (earthexplorer.usgs.gov) as raster files at a spatial resolution of 30 × 30m for every 16 days. For our analysis, we averaged each spectral index over five years, from April 2013 to April 2018, while omitting scenes that were dominated by cloud artefacts.

We extracted raster data of surface elevation from the global digital elevation model (DEM) generated using measurements from the Advanced Space-borne Thermal Emission and Reflection Radiometer (ASTER) (***Tachikawa et al., 2011***). These data are freely available for download from the USGS Earth Explorer. Using a flow accumulation map derived from the DEM, a river network map was generated and used to calculate and store as raster images the distance to small rivers and large rivers (henceforth, DSR and DLR, respectively).

### Geostatistical Analysis

The number of mosquitoes trapped by Suna traps can be used to estimate HBR, as these traps primarily target host-seeking mosquitoes. Hence, we first estimated HBR and the *P. falciparum* sporozoite rate (PfSR). We then estimated PfEIR as the product of these two quantities.

We carried out separate analyses for *A. arabiensis* and *A. funestus s*.*s*., using explanatory variables and random effects structures that we found to be suitable for each species. Details of the variable selection process and the final sets of explanatory variables for each of the models later described in this section are given in ***Appendix 1***. The correlation structures adopted for the geostatistical models were informed by the variogram-based algorithm described in (***Giorgi et al., 2018***). The geostatistical models for the HBR and PfPR data described below were fitted us-ing PrevMap (***Giorgi and Diggle, 2016***), freely available from the Comprehensive R Archive Network (CRAN, www.r-project.org). The PfSR models were fitted using the glmm package, also available on CRAN.

#### Human biting rate

Let *Y* (*x*_*i*_, *t*_*i*_), *i* = 1, …, *M*, where *M* = 2432 is the total number of households, denote counts of mosquitoes trapped at location *x*_*i*_ in month *t*_*i*_ ∈ {1, …, 38}, where *t*_*i*_ = 1 denotes April 2015. We modelled the *Y* (*x*_*i*_, *t*_*i*_) using Poisson mixed models expressed by the following linear predictor

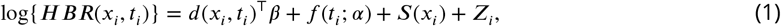

where: *d*(*x*_*i*_, *t*_*i*_) is a vector of spatiotemporal explanatory variables with associated regression coefficients ; the *f* (*t*_*i*_; *α*) is a linear combination of several functions of time, including sines, cosines and splines, with an associated vector of regression parameters *α*, accounting for trends and seasonal patterns; the *Z*_*i*_ are independent and identically distributed Gaussian random variables with variance *τ*^2^; *S*(*x*) is a zero-mean stationary and isotropic Gaussian process with variance *σ*^2^ and exponential correlation function *ρ*(*u*) = exp(−*u*/*ϕ*), where *ϕ* regulates the pace at which the spatial correlation decays for increasing distance *u* between any two locations. We allow the explanatory variables *d*(*x*_*i*_, *t*_*i*_) and *f* (*t*_*i*_; *α*) to differ between different mosquito species since different species may respond differently to environmental changes.

#### Plasmodium falciparum sporozoite rate

Let *Y* ^*^(*x*_*i*_, *t*_*i*_) be the number of mosquitoes that tested positive for the presence of *P. falciparum* sporozoites. We assumed that the *Y* ^*^(*x*_*i*_, *t*_*i*_) follow a Binomial mixed model with number of trials *N*^*^(*x*_*i*_, *t*_*i*_), i.e. the total number of successfully tested mosquitoes, and probability of testing positive *PfSR* (*x*_*i*_, *t*_*i*_). We model the latter as a logit-linear regression given by

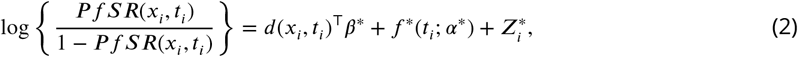

where each term in ***Equation 2*** has an analogous interpretation to those of ***Equation 1***.

Estimating the Plasmodium falciparum entomological inoculation rate Let *PfEIR* _*f*_ (*x, t*) and *PfEIR* _*a*_(*x, t*) denote the PfEIR for *A. funestus s*.*s*. and *A. arabiensis* at a given location *x* and month *t*. We estimated each of these two as

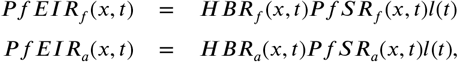

where *l*(*t*) is the number of days in month *t*. Finally, we estimated the overall PfEIR as

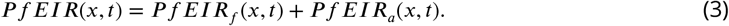

We then mapped PfEIR as in ***Equation 3*** over a 30 × 30*m* regular grid covering the whole of the study area for each month across 38 months.

#### Plasmodium falciparum prevalence

We mapped PfPR in women and in children by fitting a geostatistical model to each group. More specifically, let *I*(*x*_*i*_, *t*_*i*_) denote the number of RDT positives out of *N*_*it*_ sampled individuals at location *x*_*i*_ in month *t*_*i*_. We then assumed that the *I*(*x*_*i*_, *t*_*i*_) follow a Binomial mixed model with probability of a positive RDT result *p*(*x*_*i*_, *t*_*i*_), such that

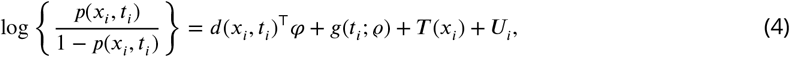

where *T* (*x*_*i*_) is a stationary and isotropic Gaussian process with exponential correlation function and *U*_*i*_ are Gaussian noise, *g*(*t*_*i*_, *ϱ*) is a linear combination of splines, and sine and cosine functions of time accounting for trends and seasonality, and *φ* and *ϱ* are vectors of regression parameters to be estimated.

#### Hotspot detection using PfEIR and PfPR

We mapped the respective predictive probabilities that PfEIR and PfPR exceeded predetermined threshold values. We then demarcated hotspots as areas where these probabilities exceeded 0.9. For PfEIR, we chose the threshold of 0.1 ib/person/month. For PfPR, we chose a threshold of 31% for children and 17% for women to correspond to the PfEIR threshold based on the best of six functional relationships between PfEIR and PfPR as described in the next section.

### Modelling the relationship between PfEIR and PfPR

In this section, we describe the statistical methods we used to model the relationship between PfEIR and PfPR. Since PfEIR may have a delayed effect on PfPR, possibly due to the time taken for *P. falciparum* to develop in the human host, we considered that current PfPR may depend on PfEIR *l* months prior. In particular, we considered *l* = 0, 1, 2. We then assumed that the number of RDT positive individuals, *I*(*x*_*i*_, *t*_*i*_), follow independent Binomial distributions such that

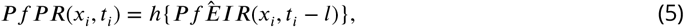

where *h*(·) is a function depending on a vector of parameters *θ* that governs the relationship between PfPR and PfEIR, and *PfÊIR* (*x*_*i*_, *t*_*i*_ − *l*) is the estimated PfEIR as in Eq (3). We considered six models, each of which provided a different specification for *h*(·).

We now describe the six models for *h*(·). Models 1 to 4 make explicit assumptions on the underlying mechanism of transmission, whereas models 5 and 6 describe the functional relationship between PfEIR and PfPR through regression methods.

#### Model 1: The susceptible-infected-susceptible (SIS) model

Let *b* be the probability that an infectious mosquito bite results in an infection, referred to as the transmission efficiency. Then, infections at (*x*_*i*_, *t*_*i*_ − *l*) are assessed to occur at a rate of *b* × *PfEIR* (*x*_*i*_, *t*_*i*_ − *l*). We assumed that each infection cleared independently over a duration 1/*r* so that the ratio *y* = *b*/*r* is the time taken to clear infection per infectious bite. We assumed that the relationship between PfEIR and PfPR holds throughout the study region. If *PfEIR* (*x, t* − *l*) is constant, the relationship between *PfEIR* (*x, t* − *l*) and *PfPR* (*x, t*) is described by the Ross Model (***Ross, 1911***)

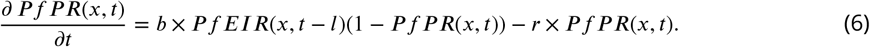

We obtained our first model as the non-zero equilibrium solution of ***Equation 6***, given by

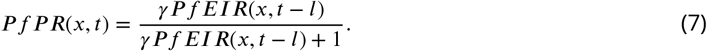

#### Model 2: The SIS model with different infection/recovery rates (D.I/R)

Model 1 assumes that women and children get infected and recover at the same rate. However, the transmission and recovery rates in children may differ from those in women. We, therefore, modified Model 1 by allowing different values of *b* and *r* for each category of people. Let *ξ*_1,*it*_ and *ξ*_2,*it*_ respectively be the proportion of children and women sampled at (*x*_*i*_, *t*_*i*_) and *y*_*k*_ = *b*_*k*_/*r*_*k*_, where *k* = 1 denotes children and *k* = 2 denotes women. The resulting Model 2 is

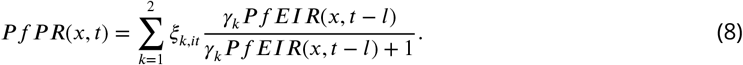

#### Model 3: The SIS model with superinfection (S.I.)

If individuals are super-infected with *P. falciparum*, then the rate at which infections clear depends on the infection rate, with clearance being faster when infection rate is low, and slower when infection rate is high. To capture this feature, we modelled infection clearance rate as *g*(*ϑ, r*) = *ϑ*/(*e*^*ϑ*/*r*^ −1), where *ϑ* = *b* × *PfEIR* (***Smith et al., 2005***; ***Walton, 1947***; ***Dletz et al., 1974***; ***Aron and May, 1982***). The resulting model for *PfPR* (*x, t*) is

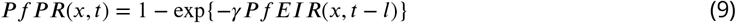

#### Model 4: The SIS model with S.I and D.I/R

Combining the assumptions of heterogeneous infection/recovery rates, as in Model 2 and superinfection, as in Model 3, we obtain Model 4,

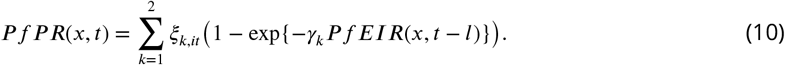

#### Model 5: The Beier model

***Beier et al***. (***1999***) assumed that the log of PfEIR is linearly related to PfPR, and fitted the regression model

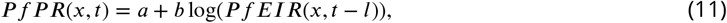

the so called “log-linear model”.

#### Model 6: The logit-linear model

The Beier model has the limitation that PfPR approaches −∞ as PfEIR goes to 0 and approaches ∞ as PfEIR goes to oo. To constrain PfPR to lie between 0 and 1, we applied the logit-link function to PfPR to give Model 6,

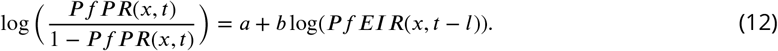

We estimated the parameters of each of the six models by maximising the log-likelihood function

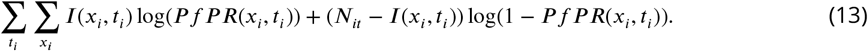

To fit each model, we first obtained 10,000 bootstrapped data sets of predicted PfEIR as in ***Equation 3*** at the set of all space-time locations sampled for the rMIS. We did this for two reasons: to obtain PfEIR data at locations (*x*_*i*_, *t*_*i*_) that were sampled for rMIS but not for the entomological surveillance; to account for the uncertainty in PfEIR. By fitting each model to each of the 10,000 datasets, we then obtained 10,000 bootstrapped samples 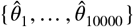 for the vector of parameter estimates 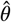 of each the six candidate models. we then summarised these samples by their mean and central 95% probability interval. We repeated this process for *l* = 0, 1, 2.

We compared the fit of the six models based on the AIC values and compared their predictive ability by the bias and root-mean-square error when each model is used to predict prevalence at all the observed space-time locations.

## Results

### rMIS and mosquito sampling

From April 2015 to May 2018, a total of 6870 traps (3439 indoors; 3431 outdoors) were placed at 2432 houses resulting in the collection of 657 female *Anopheles* mosquitoes (***Table 1***). Following PCR of the 478 *A. gambiae* s.l. collected, 92% were identified as *A. arabiensis*, 2% as *A. gambiae s*.*s*., 1% as *A. quadriannulatus*, and 5% could not be identified further. From the 179 *A. funestus* s.l. collected, 95% were identified as *A. funestus s*.*s*. by PCR, while the remaining 5% could not be identified further. The observed vector composition is therefore 71%, 27% and 2% for *A. arabiensis, A. funestus s*.*s*. and *A. gambiae s*.*s*., respectively.

**Table 1.**
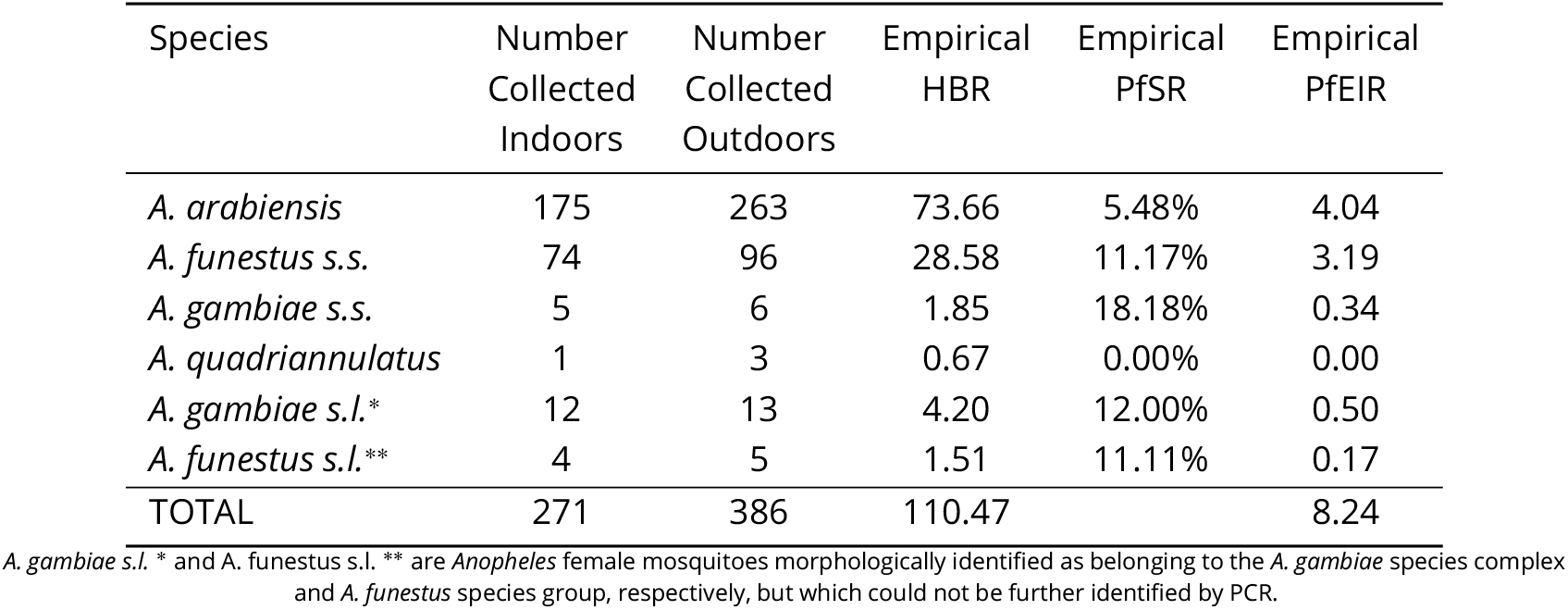
Details of *Anopheles* female mosquitoes collected. The table shows the observed numbers collected indoors and outdoors, the HBR (number collected per trap multiplied by the number of days in each of the 38 months of sampling), PfSR and PfEIR for the *Anopheles* species sampled.

Despite the relatively low abundance of *A. funestus s*.*s*. compared to *A. arabiensis*, the higher sporozoite rate of the former made the contribution of *A. funestus s*.*s*. to the total PfEIR almost equivalent to that of *A. arabiensis* (***Table 1***). The total PfEIR for the 38 months was 8.24 ib/person, equivalent to an average 2.60 ib/person/year.

Over the same 38-month period, 5685 individual *P. falciparum* RDT tests were conducted across 3096 household visits. Among the 2401 tests conducted on children aged 6 to 59 months, 25.5% were positive, while 14.3% of the 3284 tests conducted on women were positive.

### Spatiotemporal patterns of PfEIR and PfPR

We observed clear spatiotemporal heterogeneities in PfEIR, PfPR in children, and PfPR in women when mapped across the study region at a fine spatial resolution (30 x 30 m) and 1-month intervals. For convenient visualisation of the main features of the spatiotemporal maps, we have developed an interactivity web-based application to show the maps at http://chicas.lancaster-university.uk/projects/malaria_in_malawi/pfpr/. Spatially, there were differences both within and between the three focal areas. Focal Area A generally showed the lowest PfEIR and PfPR, while Focal Areas B and C showed similar, higher levels of PfEIR. Within each focal area, the spatial patterns changed from month to month, with hotspots of both PfEIR and PfPR proceeding through seasonal cycles of expansion and retraction over time. Over the 3-year study period, hotspots of PfEIR and PfPR generally disappeared during the low transmission seasons, except for residual hotspots of PfPR that persisted throughout the study period.

When summarised over the whole study region, each of PfPR and PfEIR exhibited seasonal patterns with a single annual peak. The monthly predicted PfEIRs and PfPRs were similar to the observed values (***Figure 2***). PfEIR increased from November to a peak in May and decreased to a trough in November. PfPR started increasing from December to a peak around July, after which it decreased to a trough between November/December.

**Figure 2.**
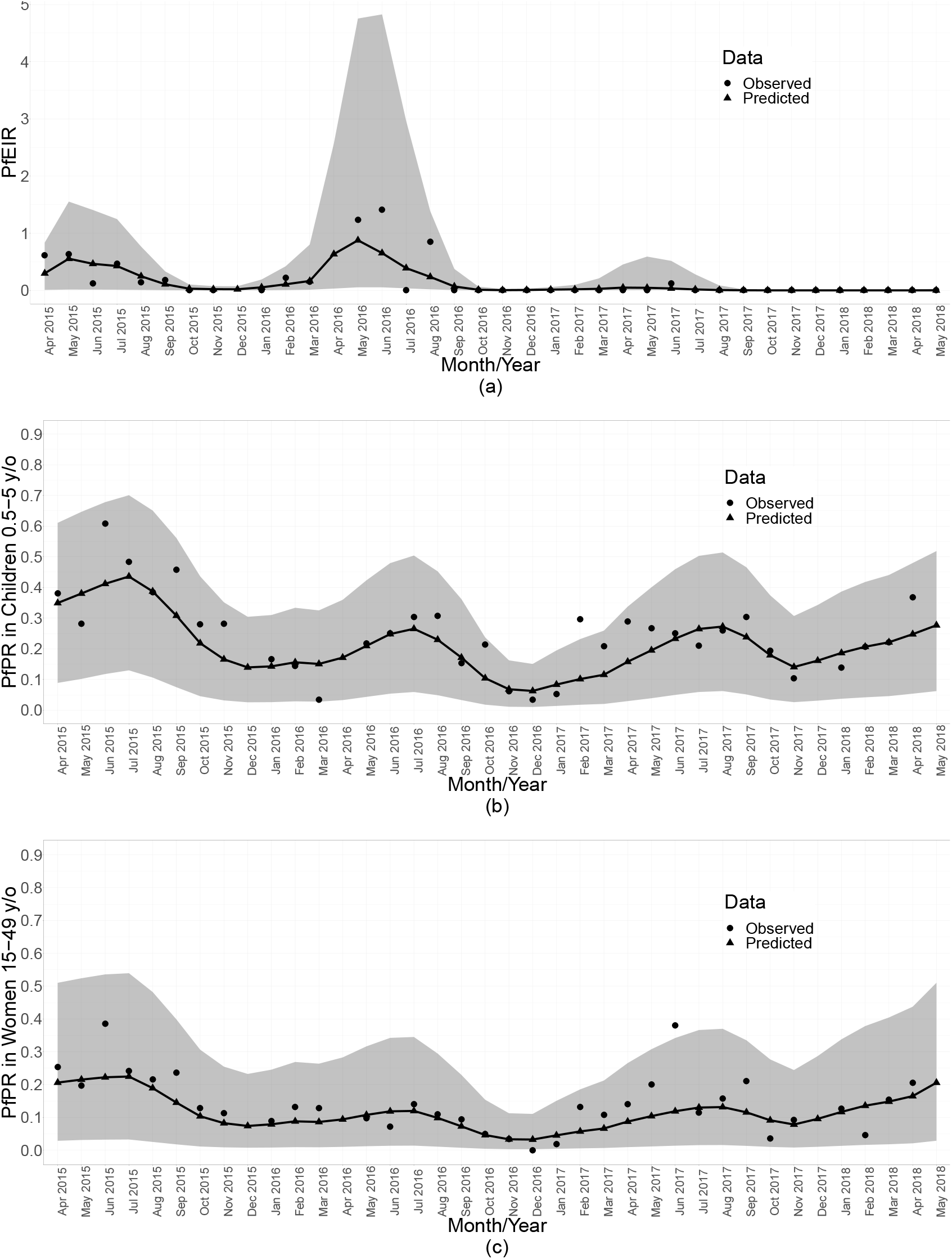
Summaries of monthly PfEIR and PfPR. The plot shows monthly median PfEIR (a), mean PfPR in children 0.5-5 y/o (b) and mean PfPR in women 15-49 y/o (c), over the study region. The round points are the observed data and the triangular points are the predictions from our models. The shaded regions represent the corresponding 95% confidence interval of the predicted values.

Three observations are clear from both the spatiotemporal maps and the monthly summarised data (***Figure 2***). First, children aged 6 to 59 months consistently had a higher level of PfPR than women throughout the study period. Second, PfPR in both groups generally decreased from the start of the study in April 2015 to December 2016, after which there was a general trend of increasing PfPR in both age groups. Finally, PfEIR was steady in the first two years of the study, followed by a general decrease after May 2016. Strikingly, the observed PfEIR was 0 between June 2017 and the end of the study, while the PfPR increased in both children and women between November 2017 and May 2018.

### The relationship between PfEIR and PfPR

Temporally, the seasonal patterns of PfEIR and PfPR within each year were nearly concurrent, with the estimated peak in PfEIR preceding that of PfPR by one month (***Figure 2***).

Spatially, PfEIR and PfPR showed broadly similar patterns. When comparing the hotspots of PfEIR and PfPR using spatiotemporal maps of exceedance probabilities, the hotspots of PfEIR and PfPR partly overlapped during the high transmission season (http://chicas.lancaster-university.uk/projects/malaria_in_malawi/pfpr/). However, there were hotspots of PfEIR that were not necessarily hotspots of PfPR and vice versa (***Figure 3***).

**Figure 3.**
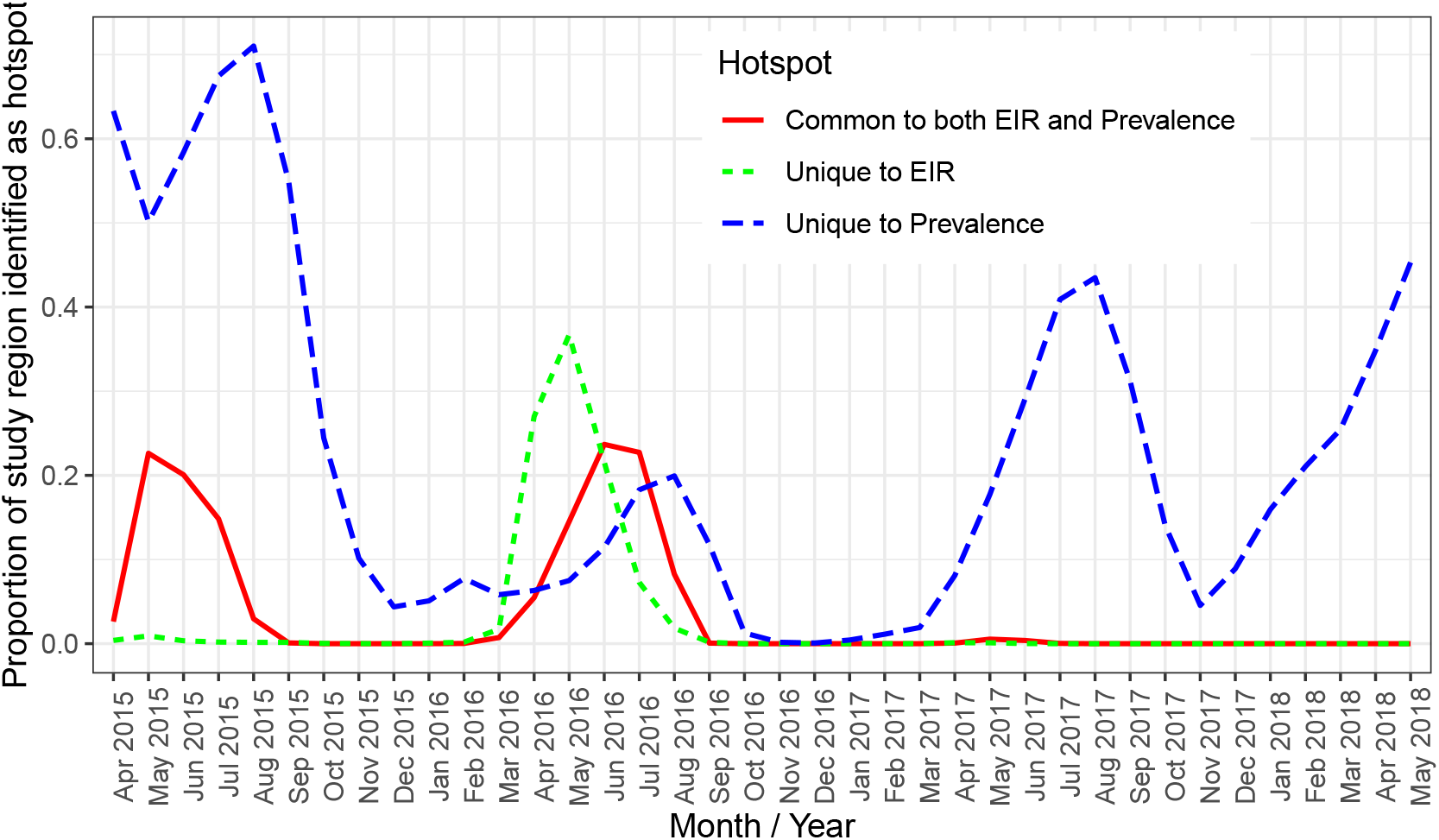
A plot of the proportion of the study region demarcated as hotspot. The solid (red) line shows hotspots identified by both PfPR and PfEIR. The long dashed (blue) line shows hotspots identified uniquely by PfPR whilst the short dashed (green) line shows hotspot uniquely identified by PfEIR.

Scatter plots of the logit of PfPR against the log of PfEIR show an approximately direct linear relationship (***Figure 4***).

**Figure 4.**
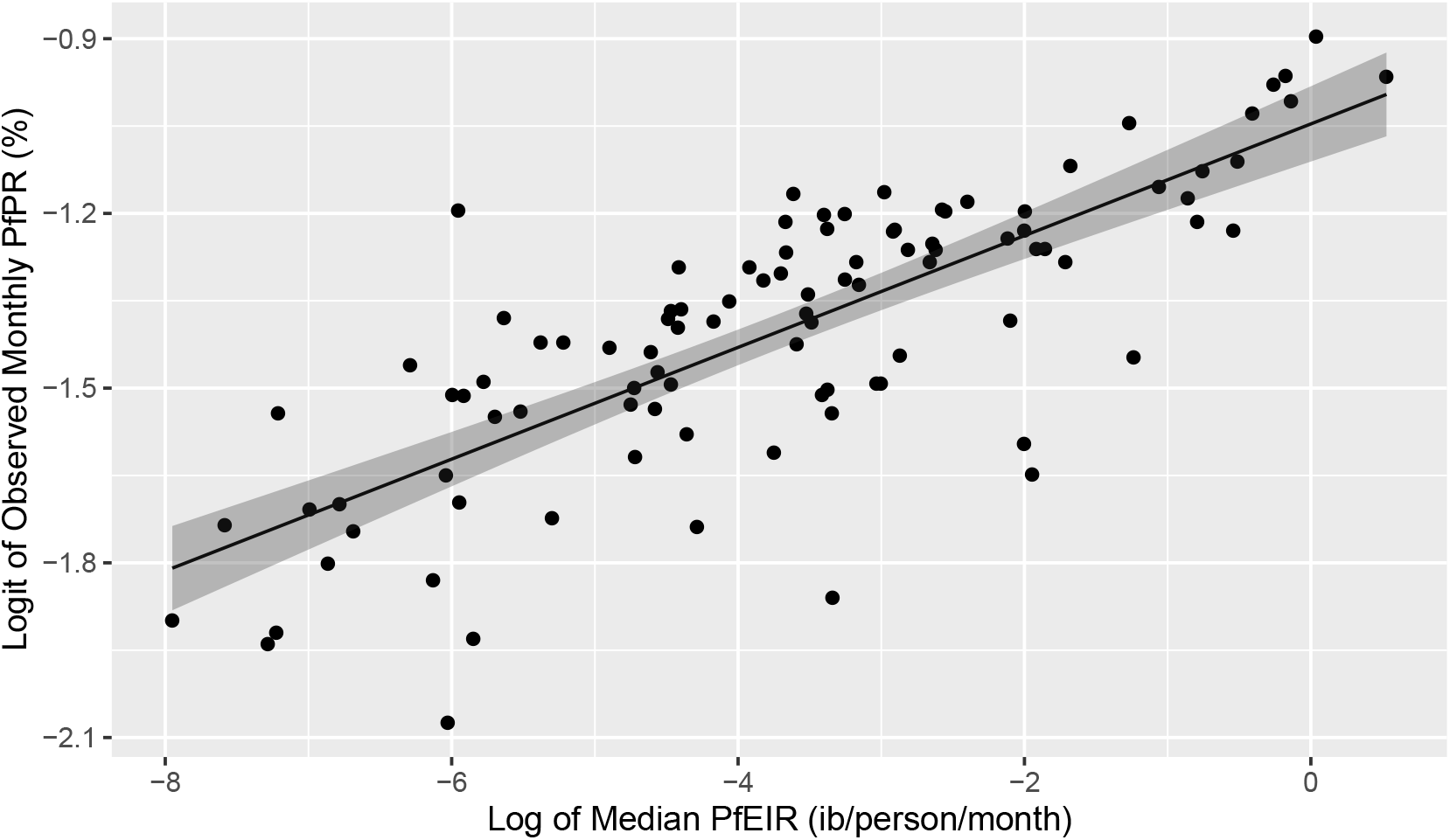
Plot of the linear relationship between the logit of PfEIR and the log of PfEIR. Each point represents a focal area and a month where there was empirical data for PfPR (n=100). PfEIR is the median (model-based predicted) PfEIR over the focal area. Prevalence is the average empirical prevalence over the focal area, with children and women put together. The shaded regions represent the corresponding 95% confidence region.

For each of the six classes of model, the model with a one-month lagged-effect gave a better fit than the corresponding models with lag zero or two (AIC differences ≥ 9). For the models with one-month lagged-effect, the empirical models (i.e. logit-linear and Beier) showed lower AIC values (i.e. better model fit, Additional Table 6 in Additional file 1) than the mechanistic models (i.e. SIS 1-4). The logit-linear model was the overall best model in terms of goodness of fit, as measured by AIC, and in terms of predictive performance, as measured by root-mean-square error and bias (***Appendix 1 Table 6***).

The fitted logit-linear model (***Figure 5***) shows that PfPR rises quickly with increasing PfEIR at very low PfEIR, followed by a flattening off or saturation. From the estimated relationship for women and children combined (***Figure 5*** (a)), a decrease in PfEIR from 1 ib/person/month to a very low PfEIR of 0.001 ib/person/month is associated with a reduction in PfPR from 27.1% to 15.8% on average (i.e., a 42.0% decrease in PfPR). Note that even when transmission, as measured by PfEIR, has been driven close to zero, PfPR remains substantial.

**Figure 5.**
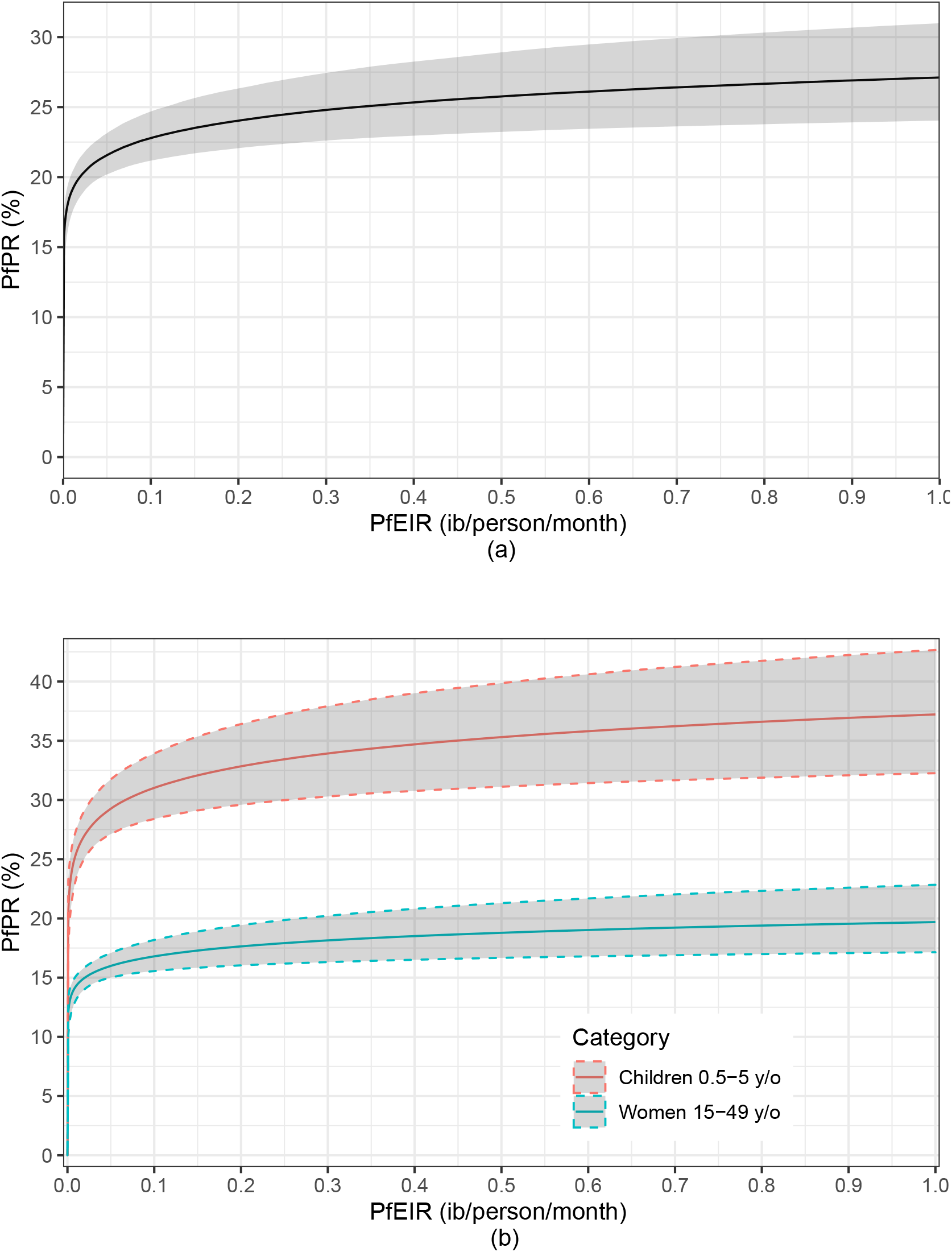
A plot of the estimated logit-linear relationship between PfPR and PfEIR. The solid lines are the estimated relationships and the shaded areas are the associated 95% confidence region for children and women combined (a) and for children and women separately (b).

An indication of differences in the PfEIR–PfPR relationship between children and women lies in the logit-linear model fitted to children and women separately (***Figure 5***(b)). The average tra-jectories of PfPR and corresponding 95% confidence intervals with varying PfEIR are distinct for women and children. PfPR in children tends to show a steeper rise with increasing PfEIR than in women. From the estimated relationship for children, a decrease in PfEIR from 1 ib/person/month to 0.001 ib/person/month is associated with a reduction in PfPR from 37.2% to 20.7% on average (i.e., a 44.5% decrease in PfPR). From the estimated relationship for women, the same decrease in PfEIR is associated with a reduction in PfPR from 19.7% to 12.1% (i.e., a 38.3% decrease in PfPR) on average. We make two observations. (1) With decreasing PfEIR, the percentage reduction in PfPR achieved in children tends to be higher than in women. (2) When transmission has been driven almost to zero, PfPR remains consistently high in children.

## Discussion

Using data from 38 months of repeated cross-sectional surveys, we have mapped the fine-scale spatiotemporal dynamics of PfEIR and PfPR in a region of Malawi with moderately intense, seasonally variable malaria parasite transmission. We found evident spatial heterogeneity in both PfEIR and PfPR, with areas of higher PfEIR and PfPR expanding and contracting over time. We also found that hotspots of PfEIR and hotspots of PfPR overlapped at times, but the amount of overlap varied over time. Finally, we showed that month-to-month variations in PfEIR over the study period are strongly associated with changes in PfPR. These findings highlight the dynamic nature of malaria parasite transmission and underscore the value of monitoring both PfEIR and PfPR at fine spatial and temporal resolutions.

Previous studies (***Beier et al., 1999***; ***Smith et al., 2005***) have demonstrated the relationship between PfEIR and PfPR using paired estimates of these metrics from several sites throughout Africa, characterised by a wide range of transmission intensities (PfEIR <1 to >500 ib/person/year). Estimates of PfEIR in our study were lower (2.6 ib/person/year, on average) so that measuring both metrics in the same geographical region, across different transmission seasons, and with a temporal resolution of one month has demonstrated that fluctuations in PfEIR over short periods are associated with predictable changes in PfPR in the same region. More specifically, our data better supported a one-month delayed effect of PfEIR on PfPR than no delayed effect or a two-month delayed effect. The one-month delayed effect is likely due to the incubation period of the parasite (***Ruan et al., 2008***) and the duration of infections (***Felger et al., 2012***). For settings with a similar range of parasite transmission intensity, our results imply that PfPR is sensitive to short-term changes in malaria parasite transmission and, therefore, can be useful for monitoring changes in the intensity of parasite transmission linked to either environmental conditions or the effects of malaria interventions. At the same time, this sensitivity to short-term changes in parasite transmission in low to moderate transmission settings suggests that single, annual, cross-sectional surveys intended to monitor inter-annual variation by aiming for a peak in PfPR are more likely to miss the actual peak than in settings with higher parasite transmission intensity (***Kang et al., 2018***). Our repeated cross-sectional sampling strategy (rolling MIS) (***Roca-Feltrer et al., 2012***; ***Kabaghe et al., 2017***) ensured that we were able to capture both short-term changes and longer-term trends in both PfEIR and PfPR. Settings where these considerations are applicable have become increasingly common over the last 20 years (***Weiss et al., 2019***), largely driven by increasing coverage of ITNs (***Bhatt et al., 2015b***) and ACTs (***Bennett et al., 2017***). Comparing 2017 with 2010, 122 million fewer people were living in areas with *PfPR* ≥ 50% (corresponding to PfEIR >15 ib/person/year), and the number of people living in settings with PfPR between 10–50% (i.e. mesoendemic (***Hay et al., 2008***)) increased to an estimated 600 million in 2017 (***Weiss et al., 2019***). In settings with higher PfEIR, PfPR likely remains relatively stable, even as PfEIR fluctuates from month to month with weather patterns (***Churcher et al., 2015***), and therefore the timing of single cross-sectional surveys is less critical.

Whereas within-village and between-village spatial heterogeneities of malaria parasite transmission are well documented across many sites (***Greenwood, 1989***; ***Thompson et al., 1997***; ***Amek et al., 2012***; ***Mwandagalirwa et al., 2017***), this is the first study of which we are aware to directly compare the fine-scale spatial patterns of PfEIR and PfPR, and these two related but distinct metrics provided a fuller picture of spatial heterogeneity in malaria parasite transmission than could have been provided by monitoring either metric in isolation. As expected, the hotspots of each metric expanded and retracted over time. However, the hotspots of PfEIR and PfPR only partially overlapped, with the most substantial amount of overlap observed during the high transmission seasons. Given the limitations of all currently available metrics of malaria parasite transmission (***Tusting et al., 2014***), our findings suggest that monitoring two transmission metrics, aligned with widely separated steps of the transmission cycle, may increase our ability to define transmission hotspots accurately. Furthermore, areas with higher transmission risk according to an entomological metric (e.g. PfEIR) than a measure of the potential transmission reservoir (e.g. PfPR) may indicate a need for increased vector control, whereas areas with lower PfEIR and higher PfPR may indicate a need for increased treatment of malaria cases (***Cohen et al., 2017***); thus, optimising the impact of control activities with minimum resources by targeting different control activities to different types of hotspots.

We found that a logit-linear model explained the PfEIR-PfPR relationship better than any of the other five model classes examined. Our results are similar to the results of ***Beier et al***. (***1999***), who assumed that the log of EIR is linearly related to PR, although our model differs from that of ***Beier et al. (1999***) in that we account for spatiotemporal heterogeneities. Our model ranking contrasts with ***Smith et al. (2005***), who favoured an SIS model analogous to our Model 4 that assumes both heterogeneous infection rates and superinfection. However, unlike the model of ***Smith et al. (2005***), we did not assume a model for age-related heterogeneities but accounted for these directly since these data were available. These differences in the overall best model between our work and that of ***Smith et al. (2005***) suggest that model performance relative to other models may be context-dependent and cautions against the use of a single model for the whole of Africa. This also highlights the importance of flexible modelling frameworks that allow accounting for spatiotemporal heterogeneity, as is the case with model-based geostatistics (***Diggle and Giorgi, 2019***).

As shown in previous studies (***Beier et al., 1999***; ***Smith et al., 2005***), our logit-linear model indicates that PfPR saturates rather than increasing at a constant rate with increasing PfEIR. This saturation in PfPR may be explained by several factors, which are not mutually exclusive. One set of factors relates to people being heterogeneously exposed to vectors (***Guelbéogo et al., 2018***) because of differences in attractiveness (***Knols et al., 1995***; ***Qiu et al., 2006***), behaviour ***Sherrard-Smith et al***. (***2019***); ***Finda et al***. (***2019***), access to ITNs (***Bhatt et al., 2015a***), housing design (***Tusting et al., 2015, 2017***), or the spatial distribution of vector habitat (***McCann et al., 2017b***), so that as PfEIR increases, it is more likely that infectious vectors are biting already infected individuals (***Smith et al., 2007b, 2010***). The second set of factors relates to inter-individual variation in acquired immunity, which in some individuals may prevent vector-inoculated sporozoites from progressing to blood-stage infection (***John et al., 2005***; ***Offeddu et al., 2017***), keep blood-stage infections at densities lower than the level of detection (***Doolan et al., 2009***) (about 50-200 parasites/µl for RDTs as used in our study), or increase the rate at which blood-stage infections are cleared (***Hviid et al., 2015***). Regardless of the reason, the saturation of PfPR has practical implications for the selection and interpretation of malaria parasite transmission metrics. When PfEIR is high, initial reductions in PfEIR will likely not be met with an immediate appreciable reduction in PfPR. Additionally, the quick rise in PfPR with increasing PfEIR at lower levels of PfEIR suggests two things, (1) that in elimination settings, a little rise in the rate of infectious bites could result in a rapid increase in parasite prevalence, making elimination extra difficult if extra efforts are not in place to avoid vector-host contacts in elimination settings; (2) that both metrics will reflect short-term changes in transmission as observed in our study.

The monthly PfEIR in our study region was 0 ib/person/month in multiple months. This may indicate that the number of infectious bites received per person during these months was below the level of detection, rather than an actual interruption of transmission during those months, especially in the first two years of the study when these periods only lasted 2-3 months. Our finding that a monthly PfEIR near or equal to zero is associated with substantial PfPR is, therefore, unsurprising given that previous studies have had similar findings when comparing annual PfEIR to PfPR (***Kabiru, 1994***; ***Mbogo et al., 1995***; ***Beier et al., 1999***; ***Smith et al., 2005***). On the other hand, we observed an increase in PfPR from about November 2017 to May 2018 while PfEIR remained at zero. It remains unclear whether this rise in PfPR was due to new infectious bites that nevertheless remained below the level of detection or to previously infected individuals with parasite densities that increased to detectable levels (***Drakeley et al., 2018***). Either way, this result shows that a rise in PfPR may be observed even when PfEIR cannot be detected by current methods, and, therefore, both interventions and monitoring need to continue for some time after PfEIR has not been detected. Our results also highlight the importance of monitoring additional metrics of parasite transmission (in addition to PfEIR) when the annual PfEIR is <10 ib/person/year, especially when expecting a reduction in transmission as in the case of testing malaria interventions. Nonetheless, when PfEIR is above the level of detection, it provides information about the vector species involved in transmission, which is critical because different mosquito species may respond differently to vector control interventions (***Ferguson et al., 2010***; ***Wilson et al., 2020***).

Prior to our study, the most recent assessment of PfEIR in this district of Malawi was from 2002, with an estimated annual PfEIR of 183 ib/person/year (***Mzilahowa et al., 2012***). The drastic reduction in annual PfEIR since then to an estimated 2.60 ib/person/year in our study is likely due, at least in part, to an increase in the use of ITNs and ACTs. Nationwide, use of ITNs by children under five years old in Malawi has increased from nil in 2000 and 14.8% in 2004 (***Mathanga et al., 2012***) to 67.8% in 2014 (***Malawi National Malaria Control Programme and ICF International, 2014***). Treatment for malaria in Malawi switched from sulfadoxine–pyrimethamine to ACT with artemether–lumefantrine in 2007 (***Mathanga et al., 2012***), and by 2014, 39.3% of children under five reporting a fever had taken ACT (***Malawi National Malaria Control Programme and ICF International, 2014***). Nationwide malaria interventions also likely impacted malaria parasite transmission over the course of our study. The most recent mass distribution of ITNs in Malawi prior to our study took place in 2012 (***World Health Organization, 2013***), with a subsequent mass ITN distribution in April 2016 that included our study site. Additionally, randomly selected villages implemented community-led larval source management, house improvement, or both as part of a randomised trial between May 2016 and May 2018 (***McCann et al., 2017a***; ***van den Berg et al., 2018***). A separate paper assesses the effects of these interventions on PfEIR and PfPR.

We observed a consistently higher PfPR in children 0.5-5 y/o than in women 15-49 y/o throughout the study region and study period, as expected. The extent of difference in PfPR between children and adults for a given region generally increases with parasite transmission intensity. However, even in mesoendemic settings (PfPR between 10–50%), it is common for PfPR in children to be appreciably higher than in adults (***Smith et al., 2007a***). This pattern is due to increasing acquired immunity with increased exposure to malaria parasites over time (***Baird, 1995***), which may decrease transmission efficiency and time to clear a *P. falciparum* infection in adults compared to children (see ***Appendix 1 Table 6*** and ***Smith et al. (2005***)). Moreover, the higher PfPR in children than adults, even at the lowest levels of transmission, suggests that children may play a more significant role in transmission, consistent with other studies (***Walldorf et al., 2015***; ***Lin Ouédraogo et al., 2015***).

One limitation of our study was the use of RDTs to estimate PfPR. RDTs can show false positives after anti-malarial drug treatment due to persistence of the antigens detected by RDTs (***Dalrymple et al., 2018***). Also, the limit of detection (usually 50-200 parasites/µl) is higher than that of expert microscopy or PCR (***Chiodini, 2014***). In modelling the relationship between PfEIR and PfPR, we did not account for the sensitivity and specificity of the RDT used to detect *P falciparum* infection. If the sensitivity *α*and specificity *β* were known, we could account for them by setting *PfPR* (*x, t*) as used in our analysis to *αPfPR* (*x, t*) + (1 − *β*)(1 − *PfPR* (*x, t*)). Thus, strictly, what we have called PfPR should be interpreted as the probability of testing positive for *P. falciparum* using RDT. However, the use of RDTs as a diagnostic test for the detection of malaria infection provides PfPR estimates that are comparable to national malaria indicator surveys.

PfPR and PfEIR are causally linked by the malaria parasite transmission cycle, which alternates between the human host and the mosquito vector. A higher rate of infectious bites received per person (i.e. EIR) increases the probability of the person becoming infected when bitten. Similarly, a higher rate of parasite infection in people (i.e. PR) increases the probability of a mosquito becoming infected after any given blood meal. Therefore, future modelling efforts may be improved by considering the EIR-PR relationship as cyclic.

## Conclusion

Measuring PfEIR and PfPR using the rolling MIS framework allowed us to assess the fine-scale spatial and temporal distributions of malaria parasite transmission over 38 months in a mesoendemic setting. The relationship between PfEIR and PfPR estimated here shows that at low transmission levels, changes in EIR are associated with rapid changes in PR, while at higher transmission levels, changes in EIR are not associated with appreciable changes in PR. Comparing hotspots of PfEIR and PfPR revealed that each metric could identify potential transmission hotspots that the other fails to capture. Our results emphasise that PfEIR and PfPR are essential, complementary metrics for monitoring short term changes in *P. falciparum* transmission intensity in mesoendemic settings, which have become increasingly common as many regions reduce transmission and shift from the highest malaria endemicity levels. Our study emphasises the need to couple vector control with identifying and treating infected individuals to drive malaria to elimination levels and to monitor both entomological and parasitaemia indices in malaria surveillance.

## Acknowledgments

This study was generously supported by Dioraphte Foundation, The Netherlands. RSM received additional support from an NIH-funded postdoctoral fellowship (T32AI007524). The content is solely the responsibility of the authors and does not necessarily represent the official views of the funders. We thank African Parks and The Hunger Project for their significant and practical contributions in facilitating the study. We are grateful to the entire Majete Malaria Project team for their tireless efforts in carrying out the study. The population of the study area is thanked for their cooperation with the study. We also thank: Alex Hiscox for advice on mosquito sampling and study design; Jeroen Spitzen for logistical assistance; and Martin Donnelley, Karl Seydel, and their respective laboratory teams for assistance in molecular identification of malaria parasites and anopheline mosquitoes.

## Appendix 1

### Procedure for building the HBR, PfSR and PfEIR models

Let *Avg*(Temp(*x*_*i*_, *t*_*i*_), *s*_1_, *s*_2_) and *Avg*(RH(*x*_*i*_, *t*_*i*_), *s*_1_, *s*_2_) respectively denote the average temperature and relative humidity taken over *s*_1_ to *s*_2_ days prior to the data collection. ***Appendix 1 Table 1*** shows the *s*_1_ and *s*_2_ values over which average temperature and relative humidity were computed. A set of these variables were selected as the best predictors each of the outcome variables based on the procedure in the next section.

We selected the best combination of fixed and random effects that best explain HBR, PfSR and PfPR using the following procedure.

1. We first built a generalized linear model in which temperature and RH are considered together with time trends and sine and cosine functions for seasonality. For *Avg*(Temp(*x*_*i*_, *t*_*i*_), *s*_1_, *s*_2_), *Avg*(RH(*x*_*i*_, *t*_*i*_), *s*_1_, *s*_2_), the choice of *s*_1_ and *s*_2_, as illustrated by ***Appendix 1 Table 1***, was based on the deviance profile of the variable involved, i.e. either temperature or RH. Piecewise-linear transformations of temperature and RH were considered based on visual inspection and epidemiological knowledge.
2. Potential confounding between seasonal sinusoids, temperature and RH were checked. Covariates that did not improve the model fit as judged by the AIC were excluded. Sincosine terms were always considered together as if they were one covariate.
3. With the current model as a basic model we include other available explanatory variables based on forward selection.
4. When no more explanatory variables significantly improve the model fit, we fit a generalized linear mixed model with a random effect for each unique space-time location.
5. We then check for the presence of residual spatial, temporal, and spatio-temporal correlations using the algorithm described in (***Giorgi et al., 2018***), and then include the random effect terms that improve the model fit.

### The selected fixed effects for the HBR, PfSR and PfPR models

We specify the set of fixed effects we selected to be in the final model for the *A. arabiensis* HBR, *A. funestus s*.*s*. HBR, PfSR, and the PfPR models. Detailed description of the terms involved in the fixed effects and the estimates of all the parameters of each model are given in S1 Tables 2 to 5.

- *A. arabiensis* human biting rate

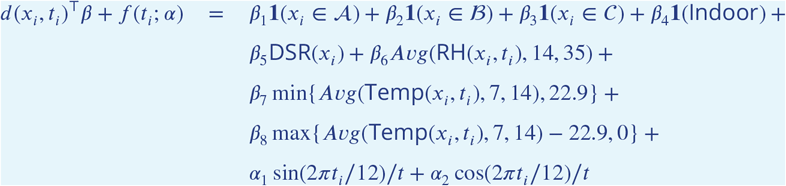
- *A. funestus s*.*s*. human biting rate

**Appendix 1 Table 1.**
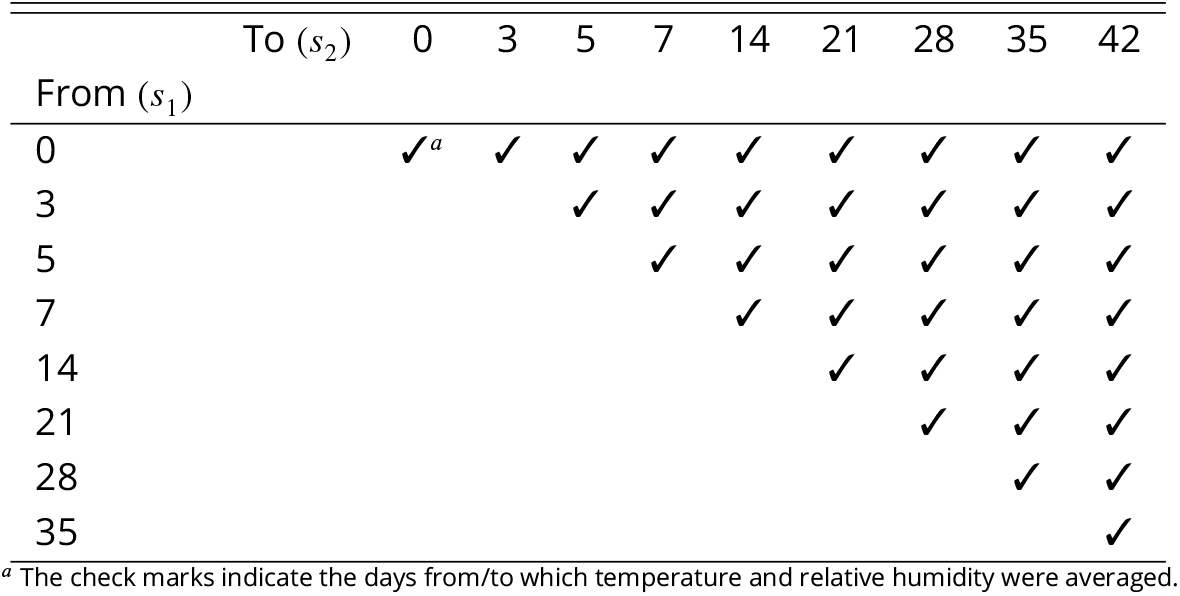
Range of days prior to data collections over which temperature and relative humidity were averaged

| | To ($s_2$) | 0 | 3 | 5 | 7 | 14 | 21 | 28 | 35 | 42 |
| --- | --- | --- | --- | --- | --- | --- | --- | --- | --- | --- |
| From ($s_1$) | | | | | | | | | | |
| 0 |  | ✓<sup>a</sup> | ✓ | ✓ | ✓ | ✓ | ✓ | ✓ | ✓ | ✓ |
| 3 |  |  |  | ✓ | ✓ | ✓ | ✓ | ✓ | ✓ | ✓ |
| 5 |  |  |  |  | ✓ | ✓ | ✓ | ✓ | ✓ | ✓ |
| 7 |  |  |  |  |  | ✓ | ✓ | ✓ | ✓ | ✓ |
| 14 |  |  |  |  |  |  | ✓ | ✓ | ✓ | ✓ |
| 21 |  |  |  |  |  |  |  | ✓ | ✓ | ✓ |
| 28 |  |  |  |  |  |  |  |  | ✓ | ✓ |
| 35 |  |  |  |  |  |  |  |  |  | ✓ |

<sup>a</sup> The check marks indicate the days from/to which temperature and relative humidity were averaged.

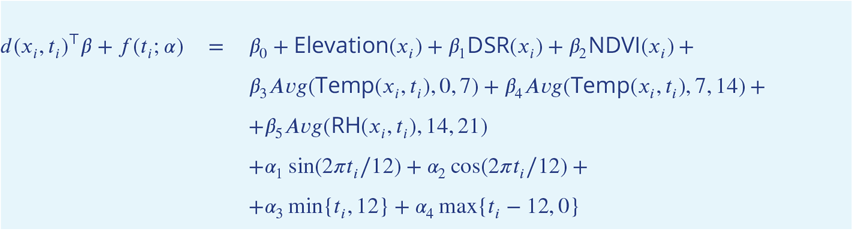
- *A. arabiensis* sporozoite rate

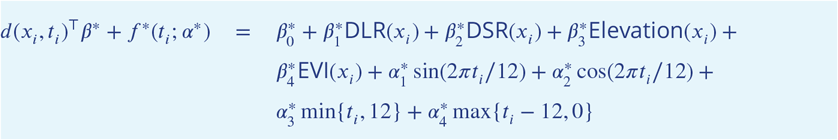
- *A. funestus s.s.* sporozoite rate

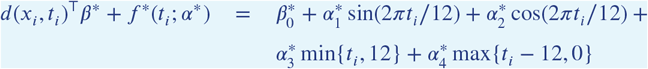
- *P. faciparum* prevalence

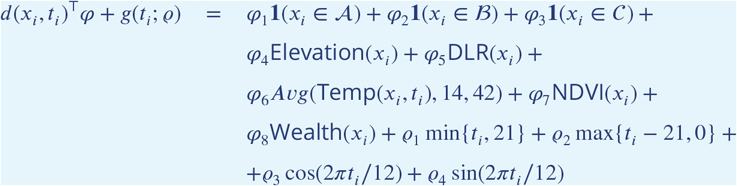

**Appendix 1 Table 2.** Regression table for the *A. arabiensis* human biting rate model

| Variable | Description | Parameter | Point Estimate |
| --- | --- | --- | --- |
| **Covariates** |  |  |  |
| $\mathbf{1}(x_i \in A)$ | A binary indicator taking the value 1 if location $x_i$ belongs to Focal Area A and 0 otherwise. | $\beta_1$ | -13.525<br>(-16.217, -10.833)<sup>a</sup> |
| $\mathbf{1}(x_i \in B)$ | A binary indicator taking the value 1 if location $x_i$ belongs to Focal Area B and 0 otherwise. | $\beta_2$ | -9.995<br>(-12.656, -7.333) |
| $\mathbf{1}(x_i \in C)$ | A binary indicator taking the value 1 if location $x_i$ belongs to Focal Area C and 0 otherwise. | $\beta_3$ | -10.848<br>(-13.514, -8.182) |
| $\mathbf{1}(\text{Indoor})$ | A binary indicator taking the value 1 if the mosquito trap was set indoors and 0 otherwise. | $\beta_4$ | 0.456<br>(0.264, 0.647) |
| $\text{DSR}(x_i)$ | Distance from location $x_i$ to the closest small river | $\beta_5$ | $0.631 \times 10^{-3}$<br>(0.143, 1.120) $\times 10^{-3}$ |
| $\text{Avg}(\text{RH}(x_i, t_i), 14, 35)$ | Average relative humidity 14 to 35 days prior to the data collection. | $\beta_6$ | 0.056<br>(0.038, 0.073) |
| $\min\{\text{Avg}(\text{Temp}(x_i, t_i), 7, 14), 22.9\}$ | The effect of temperature when temperature is below 22.9°C. | $\beta_7$ | 0.180<br>(0.072, 0.289) |
| $\max\{\text{Avg}(\text{Temp}(x_i, t_i), 7, 14) - 22.9, 0\}$ | The effect of temperature when temperature is 22.9°C or higher. | $\beta_8$ | -0.132<br>(-0.22, -0.044) |
| **Seasonality and Trends** |  |  |  |
| $\sin(2\pi t_i/12)/t$ | | $\alpha_1$ | -0.291<br>(-0.907, 0.325) |
| $\cos(2\pi t_i/12)/t$ | | $\alpha_2$ | 1.092<br>(-0.759, 2.943) |
| **Spatial Correlation** |  |  |  |
| Signal variance | | $\sigma^2$ | 4.114<br>(3.262, 5.189) |
| Scale (km) | | $\phi$ | 0.649<br>(0.492, 0.856) |
| Nugget variance | | $\tau^2$ | 0.162<br>(0.124, 0.21) |

Dependent Variable: log of *A. arabiensis* Mosquito Density<sup>a</sup> 95% confidence intervals are in brackets.

**Appendix 1 Table 3.** Regression table for the *A. funestus* human biting rate model

Dependent Variable: log of *A. arabiensis* Mosquito Density<sup>a</sup> 95% confidence intervals are in brackets. **Appendix 1 Table 3.** Regression table for the *A. funestus* human biting rate model
| Variable | Description | Parameter | Point Estimate |
| --- | --- | --- | --- |
| **Covariates** |  |  |  |
| Intercept | | $\beta_0$ | 2.523<br>(-3.209, 8.256)<sup>a</sup> |
| Elevation($x_i$) | Elevation of the location $x_i$. | $\beta_1$ | $-5.583 \times 10^{-3}$<br>(-7.896, -3.271) $\times 10^{-3}$ |
| DSR($x_i$) | Distance from location $x_i$ to the nearest small river. | $\beta_2$ | $2.993 \times 10^{-3}$<br>(2.329, 3.658) $\times 10^{-3}$ |
| NDVI($x_i$) | Normalized difference vegetation index at location $x_i$. | $\beta_3$ | 1.392<br>(-1.251, 4.035) |
| Avg(Temp($x_i, t_i$), 0, 7) | Average temperature one week prior to data collection. | $\beta_4$ | -0.154<br>(-0.279, -0.028) |
| Avg(Temp($x_i, t_i$), 7, 14) | Average temperature 7 to 14 days prior to data collection. | $\beta_5$ | -0.116<br>(-0.295, 0.064) |
| Avg(RH($x_i, t_i$), 14, 21) | Average relative humidity 14 to 21 days prior to data collection. | $\beta_6$ | -0.043<br>(-0.078, -0.008) |
| **Seasonality and Trends** |  |  |  |
| $\sin(2\pi t_i/12)$ | | $\alpha_1$ | -0.291<br>(-0.907, 0.325) |
| $\cos(2\pi t_i/12)$ | | $\alpha_2$ | 1.092<br>(-0.759, 2.943) |
| $\min\{t_i, 12\}$ | | $\alpha_3$ | -0.291<br>(-0.907, 0.325) |
| $\max\{t_i - 12, 0\}$ | | $\alpha_4$ | 1.092<br>(-0.759, 2.943) |
| **Spatial Correlation** |  |  |  |
| Signal variance | | $\sigma^2$ | 4.456<br>(3.379, 5.876) |
| Scale (km) | | $\phi$ | 0.906<br>(0.66, 1.245) |
| Nugget variance | | $\tau^2$ | 0.142<br>(0.105, 0.191) |

Dependent Variable: log of *A. funestus* Mosquito Density<sup>a</sup> 95% confidence intervals are in brackets.

**Appendix 1 Table 4.**
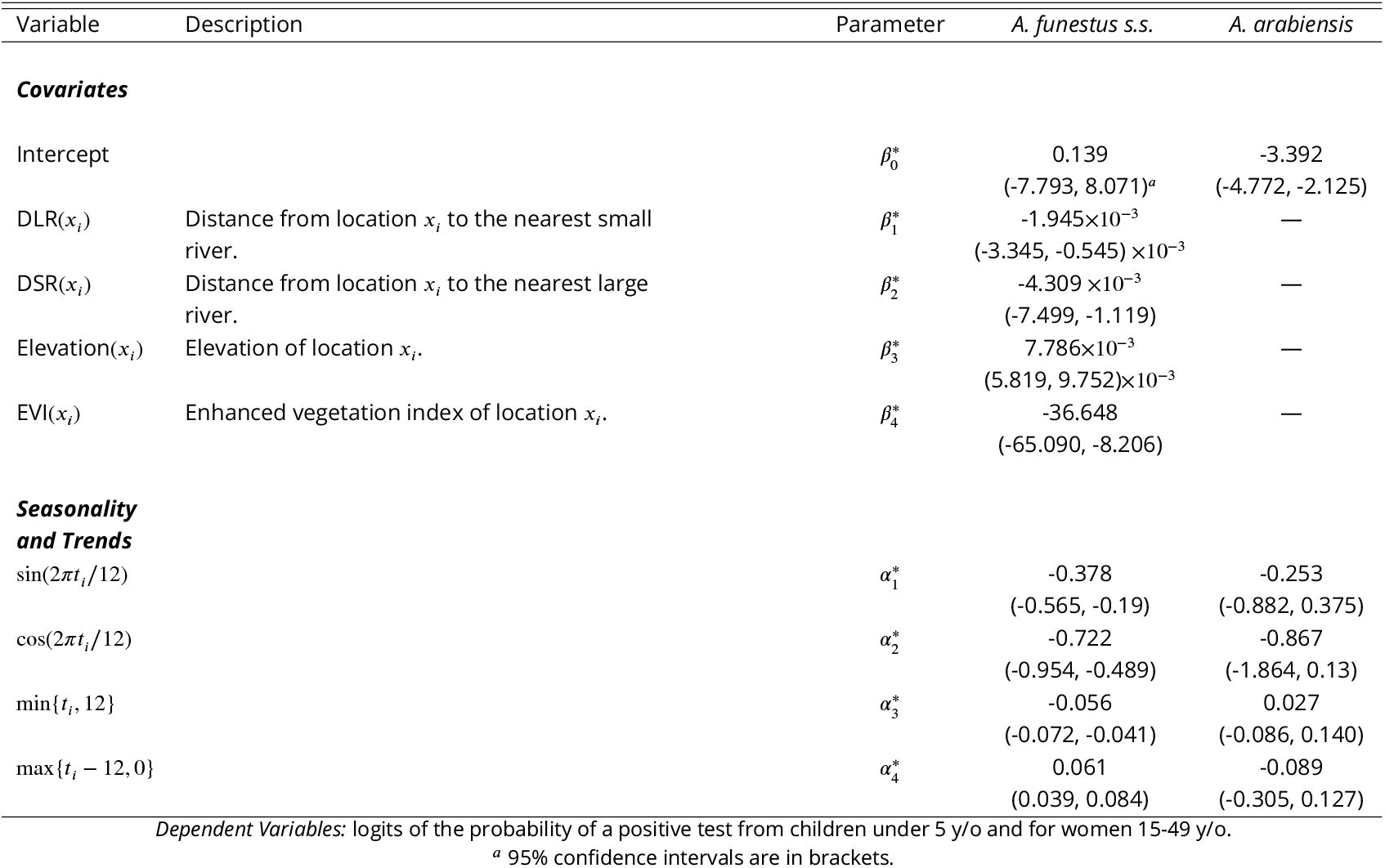
Regression table from fitting the *P. falciparum* sporozoite rate models.

**Appendix 1 Table 5.** Regression table for the *P. falciparum* parasite rate model.

| Variable | Description | Parameter | Children under 5 y/o | Women 15-49 y/o |
| --- | --- | --- | --- | --- |
| **Covariates** |  |  |  |  |
| $\mathbf{1}(x_i \in A)$ | A binary indicator taking the value 1 if $x_i$ belongs to Focal Area A and 0 otherwise. | $\varphi_1$ | 0.685<br>(-1.877, 3.247) | -0.506<br>(-3.166, 2.155) |
| $\mathbf{1}(x_i \in B)$ | A binary indicator taking the value 1 if $x_i$ belongs to Focal Area B and 0 otherwise. | $\varphi_2$ | 2.829<br>(0.41, 5.248) | 2.568<br>(0.134, 5.002) |
| $\mathbf{1}(x_i \in C)$ | A binary indicator taking the value 1 if $x_i$ belongs to Focal Area C and 0 otherwise. | $\varphi_3$ | 3.192<br>(0.806, 5.577) | 2.641<br>(0.224, 5.058) |
| Elevation($x_i$) | Elevation of the location $x_i$. | $\varphi_4$ | $5.165 \times 10^{-3}$<br>(2.322, 8.008) $\times 10^{-3}$ | $5.920 \times 10^{-3}$<br>(3.039, 8.800) $\times 10^{-3}$ |
| DLR($x_i$) | Distance from location $x_i$ to the nearest large river. | $\varphi_5$ | $-0.372 \times 10^{-3}$<br>(-0.522, -0.222) $\times 10^{-3}$ | $-0.181 \times 10^{-3}$<br>(-0.353, -0.009) $\times 10^{-3}$ |
| $\text{Avg}(\text{Temp}(x_i, t_i), 14, 42)$ | Average temperature 14 to 42 days prior to data collection. | $\varphi_6$ | -0.112<br>(-0.201, -0.023) | -0.096<br>(-0.187, -0.005) |
| NDVI($x_i$) | Normalized difference vegetation index at location $x_i$. | $\varphi_7$ | -2.424<br>(-4.703, -0.144) | -5.556<br>(-7.63, -3.482) |
| Wealth($x_i$) | Wealth index of the $i$-th household. | $\varphi_8$ | -0.212<br>(-0.283, -0.141) | -0.159<br>(-0.215, -0.102) |
| **Seasonality and Trends** |  |  |  |  |
| $\min\{t_i, 21\}$ | | $\varrho_1$ | -0.079<br>(-0.098, -0.06) | -0.079<br>(-0.1, -0.059) |
| $\max\{t_i - 21, 0\}$ | | $\varrho_2$ | 0.072<br>(0.042, 0.102) | 0.086<br>(0.056, 0.117) |
| $\cos(2\pi t_i/12)$ | | $\varrho_3$ | -0.045<br>(-0.265, 0.175) | 0.101<br>(-0.123, 0.324) |
| $\sin(2\pi t_i/12)$ | | $\varrho_4$ | 0.209<br>(-0.138, 0.556) | 0.175<br>(-0.173, 0.523) |
| **Spatial Correlation** |  |  |  |  |
| Signal variance | | $\sigma^2$ | 0.347<br>(0.222, 0.542) | 0.602<br>(0.416, 0.872) |
| Scale (km) | | $\phi$ | 1.175<br>(0.617, 2.238) | 1.055<br>(0.631, 1.765) |
| Nugget variance | | $\tau^2$ | 1.546<br>(0.956, 2.500) | 1.368<br>(0.932, 2.007) |

Dependent Variables: logits of the probability of a positive test from children under 5 y/o and for women 15-49 y/o. <sup>a</sup> 95% confidence intervals are in brackets.

**Appendix 1 Table 6.**
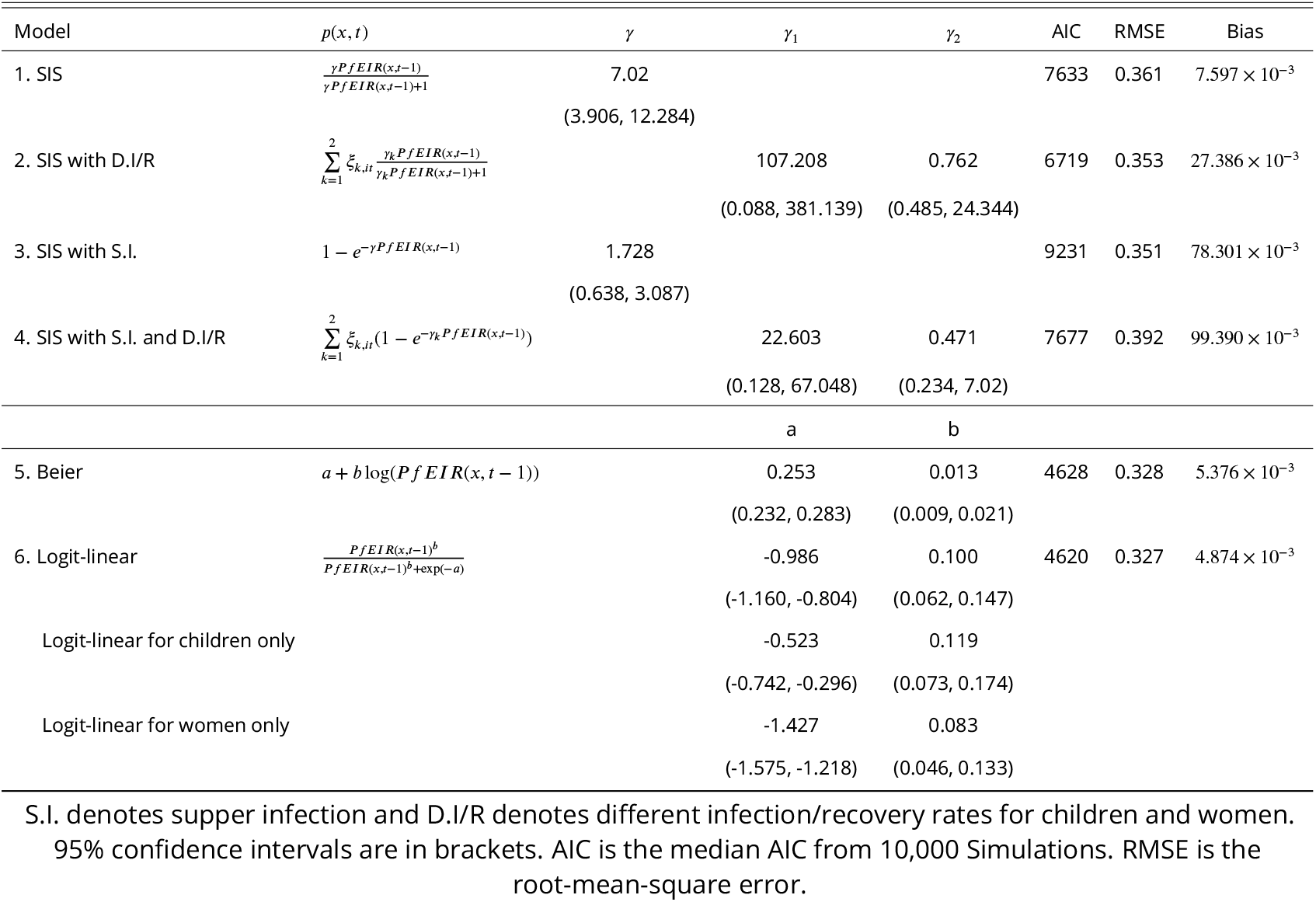
Parameter estimates from the models for the relationship between PfEIR and PfPR. The models’ goodness of fit are assessed by the AIC and their predictive abilities by the root-mean-square error (RMSE) and bias.

## References

Amek N, Bayoh N, Hamel M, Lindblade KA, Gimnig J, Laserson KF, Slutsker L, Smith T, Vounatsou P. Spatiotemporal modeling of sparse geostatistical malaria sporozoite rate data using a zero inflated binomial model. Spatial and spatio-temporal epidemiology. 2011; 2(4):283–290.

Amek N, Bayoh N, Hamel M, Lindblade KA, Gimnig JE, Odhiambo F, Laserson KF, Slutsker L, Smith T, Vounatsou P. Spatial and temporal dynamics of malaria transmission in rural Western Kenya. Parasites & vectors. 2012; 5(1):86.

Aron JL, May RM. The population dynamics of malaria. In: The population dynamics of infectious diseases: theory and applications Springer; 1982.p. 139–179.

Baird JK. Host age as a determinant of naturally acquired immunity to Plasmodium falciparum. Parasitology today. 1995; 11(3):105–111.

Bass C, Nikou D, Blagborough AM, Vontas J, Sinden RE, Williamson MS, Field LM. PCR-based detection of Plasmodium in Anopheles mosquitoes: a comparison of a new high-throughput assay with existing methods. Malaria journal. 2008; 7(1):177.

Beier JC, Killeen GF, Githure JI. Short report: entomologic inoculation rates and Plasmodium falciparum malaria prevalence in Africa. The American Journal of Tropical Medicine and Hygiene. 1999; 61(1):109–113.

Bennett A, Bisanzio D, Yukich JO, Mappin B, Fergus CA, Lynch M, Cibulskis RE, Bhatt S, Weiss DJ, Cameron E, Gething PW, Eisele TP. Population coverage of artemisinin-based combination treatment in children younger than 5 years with fever and Plasmodium falciparum infection in Africa, 2003-2015: a modelling study using data from national surveys. The Lancet Global Health. 2017 Apr; 5(4):e418–e427.

van den Berg H, Van Vugt M, Kabaghe AN, Nkalapa M, Kaotcha R, Truwah Z, Malenga T, Kadama A, Banda S, Tizifa T, Gowelo S, Mburu MM, Phiri KS, Takken W, McCann RS. Community-based malaria control in southern Malawi: a description of experimental interventions of community workshops, house improvement and larval source management. Malaria Journal. 2018 Jul; 17(1):1–12.

Bhatt S, Weiss DJ, Mappin B, Dalrymple U, Cameron E, Bisanzio D, Smith DL, Moyes CL, Tatem AJ, Lynch M, et al. Coverage and system efficiencies of insecticide-treated nets in Africa from 2000 to 2017. Elife. 2015; 4:e09672.

Bhatt S, Weiss D, Cameron E, Bisanzio D, Mappin B, Dalrymple U, Battle K, Moyes C, Henry A, Eckhoff P, et al. The effect of malaria control on Plasmodium falciparum in Africa between 2000 and 2015. Nature. 2015; 526(7572):207.

Bousema T, Griffin JT, Sauerwein RW, Smith DL, Churcher TS, Takken W, Ghani A, Drakeley C, Gosling R. Hitting Hotspots: Spatial Targeting of Malaria for Control and Elimination. PLoS Medicine. 2012 Jan; 9(1):e1001165.

Bousema T, Okell L, Felger I, Drakeley C. Asymptomatic malaria infections: detectability, transmissibility and public health relevance. Nature Reviews Microbiology. 2014; 12(12):833–840.

Bousema T, Stresman G, Baidjoe AY, Bradley J, Knight P, Stone W, Osoti V, Makori E, Owaga C, Odongo W, China P, Shagari S, Doumbo OK, Sauerwein RW, Kariuki S, Drakeley C, Stevenson J, Cox J. The Impact of Hotspot-Targeted Interventions on Malaria Transmission in Rachuonyo South District in the Western Kenyan Highlands: A Cluster-Randomized Controlled Trial. PLoS Medicine. 2016 Apr; 13(4):e1001993–25.

Carter R, Mendis KN, Roberts D. Spatial targeting of interventions against malaria. Bulletin of the World Health Organization. 2000 Jan; 78(12):1401–1411.

Chiodini PL. Malaria diagnostics: now and the future. Parasitology. 2014; 141(14):1873–1879.

Chipeta M, Terlouw D, Phiri K, Diggle P. Inhibitory geostatistical designs for spatial prediction taking account of uncertain covariance structure. Environmetrics. 2017; 28(1):e2425.

Chipeta MG, Terlouw DJ, Phiri KS, Diggle PJ. Adaptive geostatistical design and analysis for prevalence surveys. Spatial Statistics. 2016; 15:70–84.

Churcher TS, Trape JF, Cohuet A. Human-to-mosquito transmission efficiency increases as malaria is controlled. Nature Communications. 2015 Jan; 6(1):6054.

Ciota AT, Matacchiero AC, Kilpatrick AM, Kramer LD. The effect of temperature on life history traits of Culex mosquitoes. Journal of medical entomology. 2014; 51(1):55–62.

Cohen JM, Menach A, Pothin E, Eisele TP, Gething PW, Eckhoff PA, Moonen B, Schapira A, Smith DL. Mapping multiple components of malaria risk for improved targeting of elimination interventions. Malaria Journal. 2017 Nov; p. 1–12.

Craig MH, Snow R, le Sueur D. A climate-based distribution model of malaria transmission in sub-Saharan Africa. Parasitology today. 1999; 15(3):105–111.

Dalrymple U, Arambepola R, Gething PW, Cameron E. How long do rapid diagnostic tests remain positive after anti-malarial treatment? Malaria journal. 2018; 17(1):228.

Diggle PJ, Giorgi E. Model-based geostatistics for global public health: methods and applications. CRC Press; 2019.

Dletz K, et al. A malaria model tested in the African savanna. Bulletin of the World Health Organization. 1974; 50:347–357.

Doolan DL, Dobaño C, Baird JK. Acquired immunity to malaria. Clinical microbiology reviews. 2009; 22(1):13–36.

Drakeley C, Gonçalves B, Okell L, Slater H. Understanding the importance of asymptomatic and low-density infections for malaria elimination. Towards Malaria Elimination-A Leap Forward. 2018;.

Felger I, Maire M, Bretscher MT, Falk N, Tiaden A, Sama W, Beck HP, Owusu-Agyei S, Smith TA. The dynamics of natural Plasmodium falciparum infections. PloS one. 2012; 7(9).

Ferguson HM, Dornhaus A, Beeche A, Borgemeister C, Gottlieb M, Mulla MS, Gimnig JE, Fish D, Killeen GF. Ecology: a prerequisite for malaria elimination and eradication. PLoS Medicine. 2010 Aug; 7(8):e1000303.

Finda MF, Moshi IR, Monroe A, Limwagu AJ, Nyoni AP, Swai JK, Ngowo HS, Minja EG, Toe LP, Kaindoa EW, et al. Linking human behaviours and malaria vector biting risk in south-eastern Tanzania. PloS one. 2019; 14(6).

Gillies M, Coetzee M. A Supplement to the Anophelinae of Africa South of the Sahara. Publ S Afr Inst Med Res. 1987; 55:1–143.

Giorgi E, Diggle P, Prevmap: an R package for prevalence mapping. Submitted; 2016.

Giorgi E, Diggle PJ, Snow RW, Noor AM. Geostatistical Methods for Disease Mapping and Visualisation Using Data from Spatio-temporally Referenced Prevalence Surveys. International Statistical Review. 2018;.

Greenwood B. The microepidemiology of malaria and its importance to malaria control. Transactions of the Royal Society of Tropical Medicine and hygiene. 1989; 83:25–29.

Guelbéogo WM, Gonçalves BP, Grignard L, Bradley J, Serme SS, Hellewell J, Lanke K, Zongo S, Sepúlveda N, Soulama I, et al. Variation in natural exposure to anopheles mosquitoes and its effects on malaria transmission. Elife. 2018; 7:e32625.

Hay SI, Smith DL, Snow RW. Measuring malaria endemicity from intense to interrupted transmission. The Lancet Infectious Diseases. 2008 Jun; 8(6):369–378.

Hiscox A, Otieno B, Kibet A, Mweresa CK, Omusula P, Geier M, Rose A, Mukabana WR, Takken W. Development and optimization of the Suna trap as a tool for mosquito monitoring and control. Malaria journal. 2014; 13(1):257.

Hviid L, Barfod L, Fowkes FJ. Trying to remember: immunological B cell memory to malaria. Trends in parasitology. 2015; 31(3):89–94.

John CC, Moormann AM, Pregibon DC, Sumba PO, McHugh MM, Narum DL, Lanar DE, Schluchter MD, Kazura JW. Correlation of high levels of antibodies to multiple pre-erythrocytic Plasmodium falciparum antigens and protection from infection. The American journal of tropical medicine and hygiene. 2005; 73(1):222–228.

Joshua MK, Ngongondo C, Monjerezi M, Chipungu F, Liwenga E, Majule AE, Stathers T, Lamboll R. Climate change in semi-arid Malawi: Perceptions, adaptation strategies and water governance. Jàmbá: Journal of Disaster Risk Studies. 2016; 8(3):1–10.

Kabaghe AN, Chipeta MG, McCann RS, Phiri KS, Van Vugt M, Takken W, Diggle P, Terlouw AD. Adaptive geo-statistical sampling enables efficient identification of malaria hotspots in repeated cross-sectional surveys in rural Malawi. PLoS One. 2017; 12(2).

Kabiru E. Sporozoite challenge and transmission patterns as determinants of occurrence of severe malaria in residents of Kilifi district, Kenya. Nairobi: University of Nairobi. 1994;.

Kang SY, Battle KE, Gibson HS, Ratsimbasoa A, Randrianarivelojosia M, Ramboarina S, Zimmerman PA, Weiss DJ, Cameron E, Gething PW, et al. Spatio-temporal mapping of Madagascar’s Malaria Indicator Survey results to assess Plasmodium falciparum endemicity trends between 2011 and 2016. BMC medicine. 2018; 16(1):71.

Killeen GF, McKenzie FE, Foy BD, Schieffelin C, Billingsley PF, Beier JC. A simplified model for predicting malaria entomologic inoculation rates based on entomologic and parasitologic parameters relevant to control. American Journal of Tropical Medicine and Hygiene. 2000 May; 62(5):535–544.

Knols BG, de Jong R, Takken W. Differential attractiveness of isolated humans to mosquitoes in Tanzania. Transactions of the Royal Society of Tropical Medicine and Hygiene. 1995 Nov; 89(6):604–606.

Koekemoer L, Kamau L, Hunt R, Coetzee M. A cocktail polymerase chain reaction assay to identify members of the Anopheles funestus (Diptera: Culicidae) group. The American journal of tropical medicine and hygiene. 2002; 66(6):804–811.

Lin Ouédraogo A, Gonçalves BP, Gnémé A, Wenger EA, Guelbeogo MW, Ouédraogo A, Gerardin J, Bever CA, Lyons H, Pitroipa X, et al. Dynamics of the human infectious reservoir for malaria determined by mosquito feeding assays and ultrasensitive malaria diagnosis in Burkina Faso. The Journal of infectious diseases. 2015; 213(1):90–99.

Loetti V, Schweigmann N, Burroni N. Development rates, larval survivorship and wing length of Culex pipiens (Diptera: Culicidae) at constant temperatures. Journal of Natural history. 2011; 45(35-36):2203–2213.

Madder D, Surgeoner G, Helson B. Number of generations, egg production, and developmental time of Culex pipiens and Culex restuans (Diptera: Culicidae) in southern Ontario. Journal of medical entomology. 1983; 20(3):275–287.

Malawi National Malaria Control Programme, ICF International, Malawi Malaria Indicator Survey 2014. Li-longwe, Malawi, and Rockville, Maryland, USA: Malawi National Malaria Control Programme and ICF International; 2014.

Mathanga DP, Walker ED, Wilson ML, Ali D, Taylor TE, Laufer MK. Malaria control in Malawi: current status and directions for the future. Acta tropica. 2012; 121(3):212–217.

Mbogo CN, Snow RW, Khamala CP, Kabiru EW, Ouma JH, Githure JI, Marsh K, Beier JC. Relationships between Plasmodium falciparum transmission by vector populations and the incidence of severe disease at nine sites on the Kenyan coast. The American journal of tropical medicine and hygiene. 1995; 52(3):201–206.

Mburu MM, Zembere K, Hiscox A, Banda J, Phiri KS, van den Berg H, Mzilahowa T, Takken W, McCann RS. Assessment of the Suna trap for sampling mosquitoes indoors and outdoors. Malaria Journal. 2019 Feb; 18(1):1–11.

McCann RS, van den Berg H, Diggle PJ, van Vugt M, Terlouw DJ, Phiri KS, Di Pasquale A, Maire N, Gowelo S, Mburu MM, et al. Assessment of the effect of larval source management and house improvement on malaria transmission when added to standard malaria control strategies in southern Malawi: study protocol for a cluster-randomised controlled trial. BMC infectious diseases. 2017; 17(1):639.

McCann RS, Messina JP, MacFarlane DW, Bayoh MN, Gimnig JE, Giorgi E, Walker ED. Explaining variation in adult Anopheles indoor resting abundance: the relative effects of larval habitat proximity and insecticide-treated bed net use. Malaria Journal. 2017 Jul; 16(1):288.

Menger DJ, van Loon JJA, Takken W. Assessing the efficacy of candidate mosquito repellents against the background of an attractive source that mimics a human host. Medical and Veterinary Entomology. 2014 May; 28:407.

Mukabana WR, Mweresa CK, Otieno B, Omusula P, Smallegange RC, Van Loon JJ, Takken W. A novel synthetic odorant blend for trapping of malaria and other African mosquito species. Journal of chemical ecology. 2012; 38(3):235–244.

Mwandagalirwa MK, Levitz L, Thwai KL, Parr JB, Goel V, Janko M, Tshefu A, Emch M, Meshnick SR, Carrel M. Individual and household characteristics of persons with Plasmodium falciparum malaria in sites with varying endemicities in Kinshasa Province, Democratic Republic of the Congo. Malaria journal. 2017; 16(1):456.

Mzilahowa T, Hastings IM, Molyneux ME, McCall PJ. Entomological indices of malaria transmission in Chikhwawa district, Southern Malawi. Malaria journal. 2012; 11(1):380.

Offeddu V, Olotu A, Osier F, Marsh K, Matuschewski K, Thathy V. high sporozoite antibody Titers in conjunction with Microscopically Detectable Blood infection Display signatures of Protection from clinical Malaria. Frontiers in immunology. 2017; 8:488.

Onori E, Grab B. Indicators for the forecasting of malaria epidemics. Bulletin of the World Health Organization. 1980; 58(1):91–98.

Perandin F, Manca N, Calderaro A, Piccolo G, Galati L, Ricci L, Medici M, Arcangeletti M, Snounou G, Dettori G, et al. Development of a real-time PCR assay for detection of Plasmodium falciparum, Plasmodium vivax, and Plasmodium ovale for routine clinical diagnosis. Journal of clinical microbiology. 2004; 42(3):1214–1219.

Poti KE, Sullivan DJ, Dondorp AM, Woodrow CJ. HRP2: Transforming Malaria Diagnosis, but with Caveats. Trends in Parasitology. 2020 Feb; 36(2):112–126.

Qiu YT, Smallegange RC, van Loon JJA, ter Braak CJF, Takken W. Interindividual variation in the attractiveness of human odours to the malaria mosquito Anopheles gambiae s. s. Medical and Veterinary Entomology. 2006 Sep; 20(3):280–287.

Roca-Feltrer A, Lalloo DG, Phiri K, Terlouw DJ. Rolling Malaria Indicator Surveys (rMIS): a potential district-level malaria monitoring and evaluation (M&E) tool for program managers. The American journal of tropical medicine and hygiene. 2012; 86(1):96–98.

Ross R. The prevention of malaria. John Murray; London; 1911.

Ruan S, Xiao D, Beier JC. On the delayed Ross–Macdonald model for malaria transmission. Bulletin of mathematical biology. 2008; 70(4):1098–1114.

Rumisha SF, Smith T, Abdulla S, Masanja H, Vounatsou P. Modelling heterogeneity in malaria transmission using large sparse spatio-temporal entomological data. Global health action. 2014; 7.

Scott JA, Brogdon WG, Collins FH. Identification of single specimens of the Anopheles gambiae complex by the polymerase chain reaction. The American journal of tropical medicine and hygiene. 1993; 49(4):520–529.

Sherrard-Smith E, Skarp JE, Beale AD, Fornadel C, Norris LC, Moore SJ, Mihreteab S, Charlwood JD, Bhatt S, Winskill P, et al. Mosquito feeding behavior and how it influences residual malaria transmission across Africa. Proceedings of the National Academy of Sciences. 2019; 116(30):15086–15095.

Slater HC, Ross A, Felger I, Hofmann NE, Robinson L, Cook J, Gonçalves BP, Björkman A, Ouedraogo AL, Morris U, et al. The temporal dynamics and infectiousness of subpatent Plasmodium falciparum infections in relation to parasite density. Nature communications. 2019; 10(1):1–16.

Smith DL, Drakeley CJ, Chiyaka C, Hay SI. A quantitative analysis of transmission efficiency versus intensity for malaria. Nature communications. 2010; 1:108.

Smith DL, Guerra CA, Snow RW, Hay SI. Standardizing estimates of the Plasmodium falciparum parasite rate. Malaria Journal. 2007; 6(1):131–32.

Smith DL, McKenzie FE, Snow RW, Hay SI. Revisiting the basic reproductive number for malaria and its implications for malaria control. PLoS biology. 2007; 5(3).

Smith D, Dushoff J, Snow R, Hay S. The entomological inoculation rate and Plasmodium falciparum infection in African children. Nature. 2005; 438(7067):492.

Spiers A, Mzilahowa T, Atkinson D, McCall P. The malaria vectors of the lower Shire Valley, Malawi. Malawi Medical Journal. 2002; 14(1):4–7.

Stresman G, Bousema T, Cook J. Malaria Hotspots: Is There Epidemiological Evidence for Fine-Scale Spatial Targeting of Interventions? Trends in Parasitology. 2019 Aug; p. 1–13.

Stresman GH, Giorgi E, Baidjoe A, Knight P, Odongo W, Owaga C, Shagari S, Makori E, Stevenson J, Drakeley C, Cox J, Bousema T, Diggle PJ. Impact of metric and sample size on determining malaria hotspot boundaries. Scientific Reports. 2017 Apr; 7:1–8.

Tachikawa T, Hato M, Kaku M, Iwasaki A. Characteristics of ASTER GDEM version 2. In: Geoscience and remote sensing symposium (IGARSS), 2011 IEEE international IEEE; 2011. p. 3657–3660.

The malERA Refresh Consultative Panel on Characterising the Reservoir and Measuring Transmission. malERA: An updated research agenda for characterising the reservoir and measuring transmission in malaria elimination and eradication. PLoS Medicine. 2017; 14(11):e1002452–24.

Thompson R, Begtrup K, Cuamba N, Dgedge M, Mendis C, Gamage-Mendis A, Enosse SM, Barreto J, Sinden RE, Hogh B. The Matola malaria project: a temporal and spatial study of malaria transmission and disease in a suburban area of Maputo, Mozambique. The American journal of tropical medicine and hygiene. 1997; 57(5):550–559.

Tusting LS, Bottomley C, Gibson H, Kleinschmidt I, Tatem AJ, Lindsay SW, Gething PW. Housing improvements and malaria risk in sub-Saharan Africa: a multi-country analysis of survey data. PLoS medicine. 2017; 14(2).

Tusting LS, Bousema T, Smith DL, Drakeley C. Measuring changes in Plasmodium falciparum transmission: precision, accuracy and costs of metrics. In: Advances in parasitology, vol. 84 Elsevier; 2014.p. 151–208.

Tusting LS, Ippolito MM, Willey BA, Kleinschmidt I, Dorsey G, Gosling RD, Lindsay SW. The evidence for improving housing to reduce malaria: a systematic review and meta-analysis. Malaria journal. 2015; 14(1):209.

Walldorf JA, Cohee LM, Coalson JE, Bauleni A, Nkanaunena K, Kapito-Tembo A, Seydel KB, Ali D, Mathanga D, Taylor TE, et al. School-age children are a reservoir of malaria infection in Malawi. PLoS One. 2015; 10(7):e0134061.

Walton G. On the Control of Malaria in Freetown, Sierra Leon: I.—Plasmodium Falciparum and Anopheles Gambiae in Relation to Malaria Occurring in Infants. Annals of Tropical Medicine & Parasitology. 1947; 41(3-4):380–407.

Weiss DJ, Lucas TCD, Nguyen M, Nandi AK, Bisanzio D, Battle KE, Cameron E, Twohig KA, Pfeffer DA, Rozier JA, Gibson HS, Rao PC, Casey D, Bertozzi-Villa A, Collins EL, Dalrymple U, Gray N, Harris JR, Howes RE, Kang SY, et al. Mapping the global prevalence, incidence, and mortality of Plasmodium falciparum, 2000-17: a spatial and temporal modelling study. Lancet. 2019 Jul; 394(10195):322–331.

Wilson AL, Courtenay O, Kelly-Hope LA, Scott TW, Takken W, Torr SJ, Lindsay SW. The importance of vector control for the control and elimination of vector-borne diseases. PLoS Neglected Tropical Diseases. 2020; 14(1):e0007831.

World Health Organization, Focus on Malawi. Geneva: World Health Organization; 2013.

World Health Organization. Global technical strategy for malaria 2016-2030. World Health Organization; 2015.

